# IL-9 signaling redirects CAR T cell fate toward CD8+ memory and CD4+ cycling states, enhancing anti-tumor efficacy

**DOI:** 10.1101/2025.01.30.635582

**Authors:** Sofía Castelli, Wesley V. Wilson, Ugur Uslu, Amanda Finck, Charles-Antoine Assenmacher, Sebastian J. Atoche, Mikko Siurala, Regina M. Young, Carl H. June

**Affiliations:** Center for Cellular Immunotherapies, Department of Pathology and Laboratory Medicine, University of Pennsylvania Perelman School of Medicine, Philadelphia, PA, 19104, USA; Parker Institute for Cancer Immunotherapy at University of Pennsylvania, Philadelphia, PA, 19104, USA; Comparative Pathology Core, Department of Pathobiology, School of Veterinary Medicine, University of Pennsylvania, Philadelphia, PA, 19104, USA

**Keywords:** CAR T cells, cancer immunotherapy, solid tumors, cytokines, cytokine receptors, IL-9, IL-9R, T cell memory, T cell dysfunction

## Abstract

The success of chimeric antigen receptor T cell therapies targeting solid tumors is limited by the immunosuppressive tumor microenvironment. We demonstrate that endowing CAR T cells with ectopic interleukin-9 (IL-9) signaling by co-expressing an IL-9 receptor, rewires CAR T cell fate under antigen stress to enhance anti-tumor efficacy. In preclinical solid tumor models, IL-9-signaling CAR T cells exhibit increased expansion, persistence, and tumor infiltration, resulting in superior tumor control at significantly lower doses than conventional products. Trajectory and RNA velocity analyses of single-cell RNA sequencing data reveal that IL-9 signaling alters CAR T cell differentiation under antigen stress away from dysfunction, favoring a multipotent transition toward CD8+ cell memory and effector states, and promoting a CD4+ cell proliferative state. Interrogation of transcription factor pathways indicates that IL-9-mediated activation of STAT1 and STAT4 drives the superior phenotype of IL-9-signaling CAR T cells, providing a promising therapeutic strategy for targeting solid cancers.

## INTRODUCTION

Chimeric antigen receptor (CAR) T cell therapy has revolutionized the treatment of hematologic malignancies [1, 2]. However, its efficacy against solid tumors is hindered by challenges such as limited T cell trafficking and infiltration, an immunosuppressive tumor microenvironment (TME), and on-target off-tumor toxicity [3]. Enhancing CAR T cell fitness is crucial to overcome these barriers.

Engineering CAR T cells to incorporate cytokine signaling pathways has shown promise in addressing these obstacles [4]. For example, interleukin-2 (IL-2) supports T cell proliferation and survival but can cause severe adverse effects when administered systemically due to widespread receptor expression [5–7]. To mitigate these effects, strategies such as orthogonal IL-2 receptor systems — engineered receptor-ligand pairs that minimize interactions with native cytokine pathways — are under investigation (NCT05665062). Similarly, IL-15 secretion enhances CAR T cell antitumor efficacy in preclinical studies [8, 9], but systemic administration risks immune overactivation [10]. Clinical trials (NCT05103631) have reported improved response rates but also a higher incidence of cytokine release syndrome [11], emphasizing the need for precise control mechanisms. IL-12-armored CAR T cells have shown improved cytotoxicity and TME remodeling in preclinical studies, with ongoing clinical trials testing this approach (NCT02498912, NCT06343376) [12, 13]. However, systemic IL-12 secretion induces severe toxicities, even when under inducible control [14]. While these cytokine-enhanced designs address TME challenges, constitutive cytokine secretion often exacerbates toxicity risks due to the broad expression of cytokine receptors, underscoring the need for safer strategies [12, 13].

Incorporating IL-9 signaling into CAR T cells represents a novel strategy for overcoming these barriers. IL-9, a common gamma chain (γ_c_) cytokine, exerts diverse effects on tumor immunity, including enhanced immune cell recruitment and modulation, yet remains underexplored [15–19]. Notably, unlike receptors for other γ_c_ cytokines such as IL-2 and IL-15, the IL-9 receptor (IL-9R) has minimal expression in T cells and across normal tissues (Jiang et al., 2024). This unique expression profile minimizes toxicity risks, positioning IL-9-engineered CAR T cells as a safer and potentially more effective therapeutic alternative.

Our prior work demonstrated that CAR T cells engineered with an orthogonal IL-9 receptor (o9R) could effectively enhance antitumor immunity in murine solid tumor models [20]. However, emerging evidence suggests that the native IL-9R may confer distinct advantages over its orthogonal counterpart (Jiang et al., 2024). Specifically, IL-9R-engineered T cells exhibit superior antitumor efficacy, with enhanced peripheral expansion, greater tumor infiltration, improved memory T cell formation, and stronger STAT phosphorylation compared to o9R-engineered T cells (Jiang et al., 2024). These findings suggest that the natural IL-9 receptor, rather than engineered orthogonal or mutated counterparts, better amplifies endogenous signaling pathways to drive more robust antitumor responses.

Here, we engineered CAR T cells to express an authentic IL-9R (CAR-IL9R T cells), enabling IL-9 responsiveness. Combined with intratumoral adenoviral IL-9 delivery, CAR-IL9R T cells showed improved antitumor efficacy and survival in preclinical models of pancreatic ductal adenocarcinoma (PDA) and prostate cancer. IL-9 signaling promoted enhanced CAR T cell expansion, persistence, and tumor infiltration, while preserving functionality and central memory phenotypes in an *in vitro* dysfunction model. By leveraging the intrinsic advantages of the native IL-9R, our work examines its unique contributions to human CAR T cell function, aiming to enhance the efficacy and precision of CAR T cell therapies for solid tumors.

## RESULTS

### Authentic IL-9 signaling enhances CAR T cell antitumor efficacy in a syngeneic PDA model

Equipping CAR T cells with cytokine receptors such as IL-9R, whose expression is restricted to a limited subset of human cells, offers a strategy for selectively enhancing CAR T cell activity (Jiang et al., 2024). To investigate how the introduction of authentic IL-9 signaling impacts cell extrinsic and intrinsic mechanisms governing CAR T cell function, we first employed a murine syngeneic PDA model. Mouse T cells were engineered to express a mesothelin-targeted CAR (A03 CAR) and the wildtype IL-9R (CAR-IL9R T cells) (**Figures 1A and 1B**). IL-9 treatment resulted in the dose-dependent phosphorylation of STAT1, STAT3, and STAT5, consistent with known IL-9R signaling pathways [21] (**Figure 1C**). *In vitro* functional assays revealed that IL-9-treated CAR-IL9R T cells more effectively killed mesothelin-expressing PDA7940b cells, derived from KPC (Kras^LSL.G12D/+^p53^R172H/+^) mice, than control CAR T cells (**Figure 1D**). This enhancement was associated with increased production of cytokines, including IFN-γ, TNF-α, and IL-10 (**Figure 1E**). Additionally, IL-9 treatment induced a shift toward a stem-like memory T cell phenotype (CD44-CD62L+CD95+) in CAR-IL9R T cells (**Figure 1F**), mirroring findings from our previous studies with o9R-engineered T cells [20].

**Figure 1:**
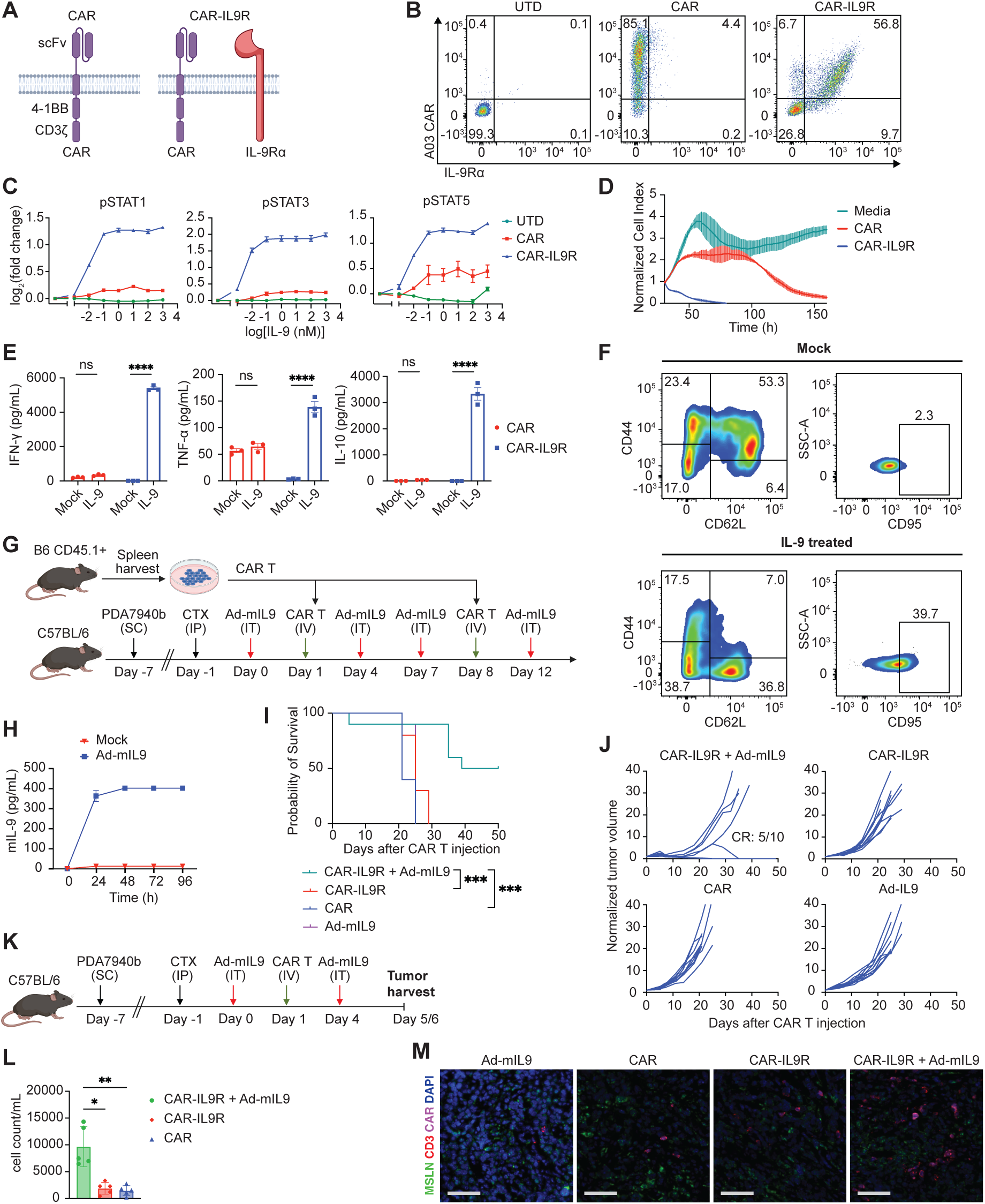
**Authentic IL-9 signaling enhances CAR T cell antitumor efficacy in a syngeneic PDA model** (A) Schematic representation of T cells transduced to express either CAR alone (CAR) or CAR and IL-9R (CAR-IL9R). (B) Flow cytometric analysis of transduction efficiency in murine untransduced (UTD), CAR, or CAR-IL9R T cells. (C) Quantification of phosphorylated STAT1 (pSTAT1), STAT3 (pSTAT3) and STAT5 (pSTAT5) in UTD, CAR, or CAR-IL9R T cells following 20-minute IL-9 stimulation. Data are shown as log fold change of mean fluorescence intensity. Error bars indicate mean ± standard error of the mean (SEM) (n = 3 biological replicates). (D) *In vitro* cytotoxicity assay of PDA7940b tumor cells co-cultured with CAR or CAR-IL9R T cells (3:1 E:T ratio), with T cells pre-incubated with IL-9 for 48 hours. Media was used as a control. Error bars indicate mean ± standard deviation (SD) (n = 3). (E) Cytokine secretion profiles of CAR and CAR-IL9R T cells, either untreated (Mock) or pre-incubated with IL-9 for 48 hours. Error bars indicate mean ± SEM (n = 3 biological replicates). (F) Representative flow cytometry plots showing surface expression of CD44, CD62L and CD95 on CAR-IL9R T cells, either untreated (Mock) or treated with IL-9 for 48 hours. (G)Experimental design schematic for the syngeneic PDA model. Ad-mIL9 dose: 10^9^ VP; CAR T cell dose: 5 × 10^6^ cells; cyclophosphamide (CTX) dose: 120 mg/kg. Administration routes: subcutaneous (SC); intraperitoneal (IP); intratumoral (IT); intravenous (IV). (H) *In vitro* murine IL-9 (mIL-9) production by PDA7940b cells treated with Ad-mIL9, measured by ELISA. Error bars indicate mean ± SD (n = 2). (I) Kaplan-Meier survival curves (n = 10 per cohort). (J) Tumor volume progression over time. CR: complete remission. (K) Experimental design schematic for tumor microenvironment analysis. Tumors were collected on day 5 post-treatment for immunohistochemistry and RNA in situ hybridization (RNA ISH), and on day 6 for flow cytometric analysis in separate experiments. (L) Quantification of infused (CD45.1+) T cells from tumor single cell suspensions analyzed via flow cytometry. Error bars indicate mean ± SD (n = 5 per cohort). (M)Representative RNA ISH photomicrographs (20x magnification; scale bar: 70μm). Illustrations were created with Biorender.com. Statistical significance was calculated by two-way ANOVA with Bonferroni correction for multiple comparisons or Kruskal-Wallis one-way ANOVA. For Kaplan-Meier survival analysis, the log-rank Mantel-Cox test was used. Statistical significance is denoted as follows: ns (not significant) *P* > 0.05; \**P* ≤ 0.05; \*\**P* ≤ 0.01; \*\*\**P* ≤ 0.001; \*\*\*\**P* ≤ 0.0001. See also Figure S1.

Next, we evaluated the influence of the TME on the antitumor efficacy of CAR-IL9R T cells *in vivo* (**Figure 1G**). C57BL/6 mice were implanted subcutaneously with PDA7940b cells. T cells harvested from CD45.1+ B6.SJL-Ptprc^a^ Pepc^b^/BoyJ mice were subjected to retroviral transduction to produce CAR-IL9R T cells. CAR and IL-9R transduction efficiency was confirmed by flow cytometry before intravenous (IV) injection into recipient mice. To provide IL-9 signaling, mice received intratumoral (IT) injections of a replication-deficient adenovirus encoding murine IL-9 (Ad-mIL9). Prior to *in vivo* testing, IL-9 production by tumor cells after Ad-mIL9 infection was validated *in vitro* (**Figure 1H**). Control groups for *in vivo* studies included mice treated with CAR-IL9R T cells alone, CAR T cells alone, or Ad-mIL9 alone. Treatment with CAR-IL9R + Ad-mIL9 significantly improved overall survival (**Figure 1I**) and reduced tumor progression (**Figure 1J**). Toxicity characterized by clinical signs of immune-effector-cell-associated neurotoxicity syndrome (ICANS) was observed in 1/10 mice treated with CAR-IL9R + Ad-mIL9. This phenomenon was observed at higher levels (5/12 mice) following treatment with CAR T cells expressing o9R and was investigated in our previous study [20]. Briefly, results suggested that toxicity was driven by CAR on-target off-tumor activity, due to the expression of mesothelin in the murine, but not healthy human meningeal layers [20, 22].

Given the marked antitumor efficacy observed with CAR-IL9R + Ad-mIL9 in this immunologically “cold” PDA model, we investigated potential changes in the TME. Tumors, spleens, and blood were collected on days 5/6 post-treatment for analysis via flow cytometry, immunohistochemistry (IHC), and RNA in situ hybridization (ISH) (**Figure 1K**). Tumors from the CAR-IL9R + Ad-mIL9 group showed significantly higher CAR T cell infiltration, as indicated by an increase in CD45.1+ (infused) T cells detected via flow cytometry (**Figure 1L**) and higher levels of CAR RNA determined by RNA ISH (**Figures 1M** and **S1A**). The spleens from this group also exhibited higher CAR T cell numbers compared with the group treated with conventional CAR T cells (**Figure S1B**, left), while no significant differences were observed in blood CAR T cell numbers (**Figure S1B**, right) or serum cytokine levels (**Figure S1C**). No significant differences in other immune cell populations within the TME were observed across treatment groups, as determined by flow cytometry (**Figure S1D**). This analysis included B cells (CD19+), NK cells (NK1.1+), dendritic cells (F4/80-CD86+CD11c+), M1 macrophages (F4/80+CD86+MHCII+), M2 macrophages (F4/80+CD86+MHCII-CD206+), endogenous T cells (CD3+CD45.1-), granulocytic MDSCs (CD11b+Ly6G+Ly6C-), and monocytic MDSCs (CD11b+Ly6G-Ly6C+). These findings were corroborated by IHC (**Figure S1E**), which showed no significant differences in B cells (PAX5+), Tregs (FoxP3+), mast cells (Tryptase+), macrophages (F4/80+), dendritic cells (CD11c+), or eosinophils (PRG2+). While there was an increase in NK cells (K1RB1C+) in the CAR-IL9R + Ad-mIL9 group compared with the CAR-IL9R group, this was not statistically significant compared to the CAR-treated group. Additionally, fewer Ly6G+ cells were observed in the CAR-IL9R + Ad-mIL9 group, possibly indicating reduced neutrophil infiltration. Overall, IL-9-signaling CAR T cells did not substantially alter cell type composition within the TME.

### Human IL-9-signaling CAR T cells exhibit enhanced potency and persistence in a xenograft PDA model

To extend our findings to a human system, we next evaluated the antitumor efficacy of human IL-9-signaling CAR T cells in a xenograft flank PDA model (**Figure 2A**). Immunodeficient NOD/scid/IL2rγ^−/−^(NSG) mice were implanted with the human mesothelin-expressing PDA cell line, AsPC-1. Primary human T cells derived from healthy donors were transduced with a human mesothelin-specific CAR construct (M5 CAR) and IL-9R to enable IL-9 signaling. CAR and IL-9R transduction efficiency was assessed via flow cytometry before IV injection into mice. To provide an IL-9 signal to CAR-IL9R T cells, an IT injection of adenovirus encoding human IL-9 (Ad-hIL9) was administered. Prior to *in vivo* testing, IL-9 production from Ad-hIL9-transduced tumor cells was confirmed *in vitro* (**Figure 2B**). A dose titration study revealed that doses as low as 0.03E6 CAR-IL9R T cells combined with Ad-hIL9 were sufficient to achieve tumor clearance (data not shown). Further survival analysis demonstrated that this low dose combination resulted in significantly enhanced antitumor efficacy compared to CAR T cells without IL-9 signaling at a 1E6 dose (**Figures 2C** and **2D**), underscoring the potency of IL-9-signaling CAR T cells.

**Figure 2:**
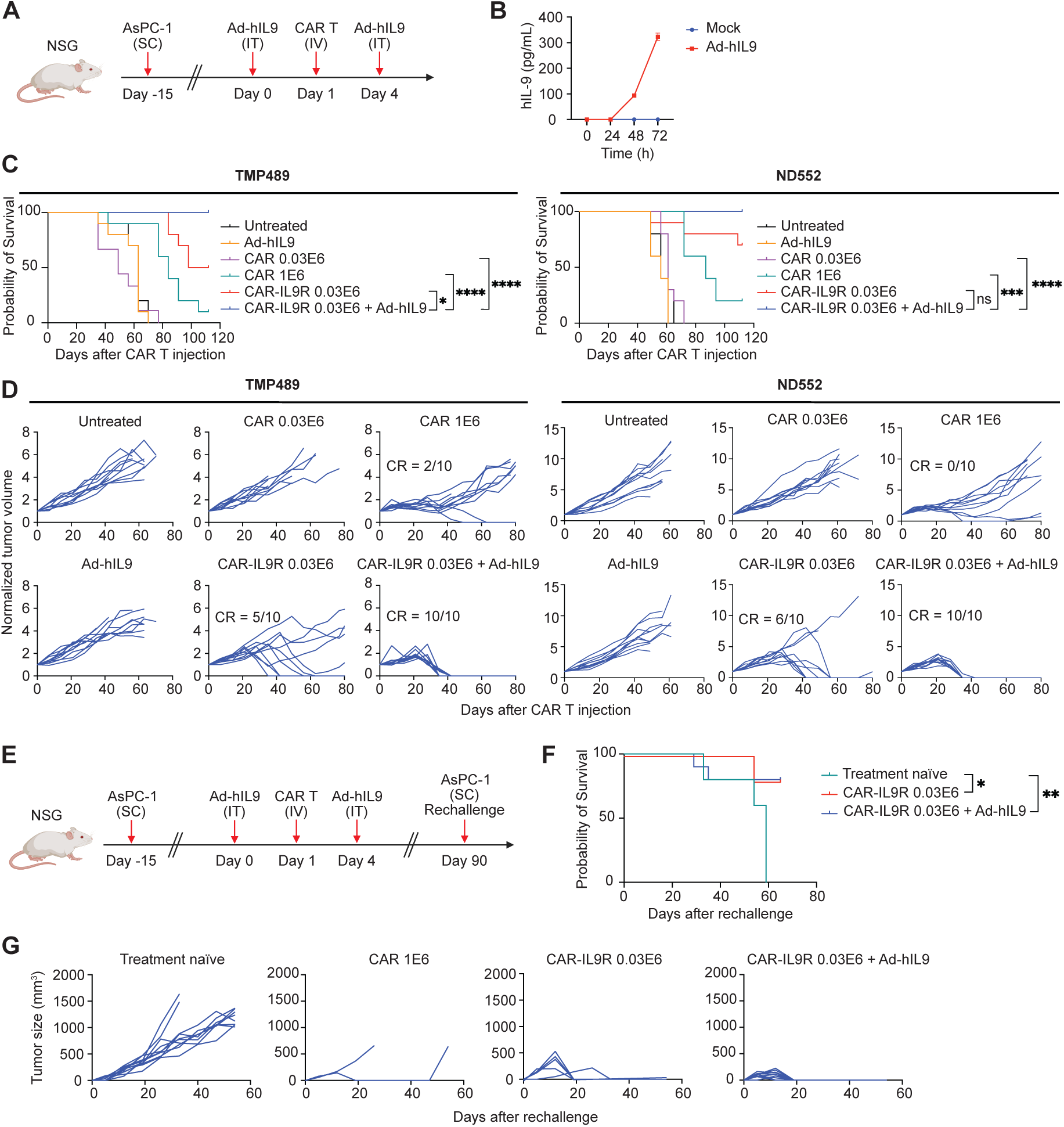
Human IL-9-signaling CAR T cells exhibit enhanced potency and persistence in a xenograft PDA model. (A) Schematic of the xenograft PDA model. Ad-hIL9 dose: 10^9^ VP. CAR T cell doses: 0.03 × 10^6^ cells (CAR 0.03E6, CAR-IL9R 0.03E6 and CAR-IL9R 0.03E6 + Ad-hIL9 cohorts) or 1 x 10^6^ cells (CAR 1E6 cohort). Administration routes: subcutaneous (SC); intratumoral (IT); intravenous (IV). (B) *In vitro* production of human IL-9 (hIL-9) by AsPC-1 cells infected with Ad-hIL9, measured by ELISA. Error bars indicate mean ± SD (n = 3). (C) Kaplan-Meier survival curves (n = 10 per cohort) from two independent experiments using CAR T cells produced from two normal donors, TMP489 (left) and ND552 (right). (D) Tumor volume progression over time. CR: complete remission. (E) Schematic of tumor re-challenge study design. (F) Kaplan-Meier survival curves for treatment-naïve mice (n=10), CAR-IL9R 0.03E6 cohort (n=5) and CAR-IL9R + Ad-hIL9 cohort (n=10). (G)Tumor volume progression over time following tumor re-challenge. Illustrations were created with Biorender.com. Statistical significance was calculated using the log-rank Mantel-Cox test. ns (not significant) *P* > 0.05; \**P* ≤ 0.05; \*\**P* ≤ 0.01; \*\*\**P* ≤ 0.001; \*\*\*\**P* ≤ 0.0001. See also Figure S2 and Table S1.

To assess the *in vivo* long-term persistence of IL-9-signaling CAR T cells, we conducted a tumor rechallenge experiment. On day 90 after treatment, AsPC-1 cells were implanted into the contralateral flank of mice that had achieved complete remission of their primary tumors (**Figure 2E**), with treatment-naïve mice serving as controls. Mice previously treated with IL-9-signaling CAR T cells rejected the newly implanted tumors and demonstrated significantly improved overall survival compared to naïve controls (**Figures 2F** and **2G**). While 2/10 mice previously treated with CAR-IL9R + Ad-hIL9 were euthanized due to signs of graft-versus-host disease (GVHD) at late timepoints, it is noteworthy that none of these mice developed tumors, highlighting the high persistence of IL-9-signaling CAR T cells.

Notably, the group receiving CAR-IL9R T cells without Ad-hIL9 also exhibited enhanced overall survival and reduced tumor progression, both in the primary tumor and upon rechallenge, as well as enhanced blood CAR T cell expansion, when compared to CAR T cells lacking IL-9R expression (**Figures 2C-2G**). Given that NSG mice have mast cells capable of producing murine IL-9, which cross-reacts with human IL-9R [23], and that T cells can also produce IL-9 [21], we hypothesized that either murine IL-9 from NSG mice or human IL-9 from the infused CAR T cells was responsible for activating IL-9R on CAR-IL9R T cells. To test our hypothesis, we measured the levels of both murine IL-9 (mIL-9) and human IL-9 (hIL-9) in the serum of treated mice. Our analysis revealed that hIL-9 was only detected in the group treated with Ad-hIL9 (**Figure S2A**, left panel), while it was undetectable in the CAR-IL9R + Ad-hIL9 group, likely due to sequestration by IL-9R-expressing CAR T cells. However, mIL-9 was detected across all treatment groups (**Figure S2A**, right panel). This led us to speculate that mIL-9 was responsible for activating IL-9R on CAR-IL9R T cells, contributing to the enhanced antitumor efficacy observed in the group treated with CAR-IL9R T cells alone. To test this, we treated mice with a neutralizing antibody targeting mIL-9 (α-mIL9) (**Figure S2B**). As expected, neutralizing mIL-9 abrogated the enhanced antitumor effects of CAR-IL9R T cells (**Figures S2C** and **S2D**). Furthermore, mice treated with CAR-IL9R T cells and α-mIL9 showed a reduced number of CAR T cells in the blood (**Figure S2E**). To further explore the potential impact of circulating IL-9 in a clinical context, we assessed hIL-9 levels in the serum of both healthy donors and patients with pancreatic cancer before and after M5 CAR T cell infusion. In our analysis, hIL-9 was undetectable in all samples (**Table S1**).

### IL-9 signaling enhances CAR T cell expansion, persistence, and tumor infiltration *in vivo*

Flow cytometry analysis on blood samples revealed that IL-9-signaling CAR T cells expanded significantly more than conventional CAR T cells and persisted for up to 10 weeks post-treatment (**Figure 3A**). This expansion was associated with increased cytokine production, suggesting enhanced effector function (**Figure 3B**).

**Figure 3:**
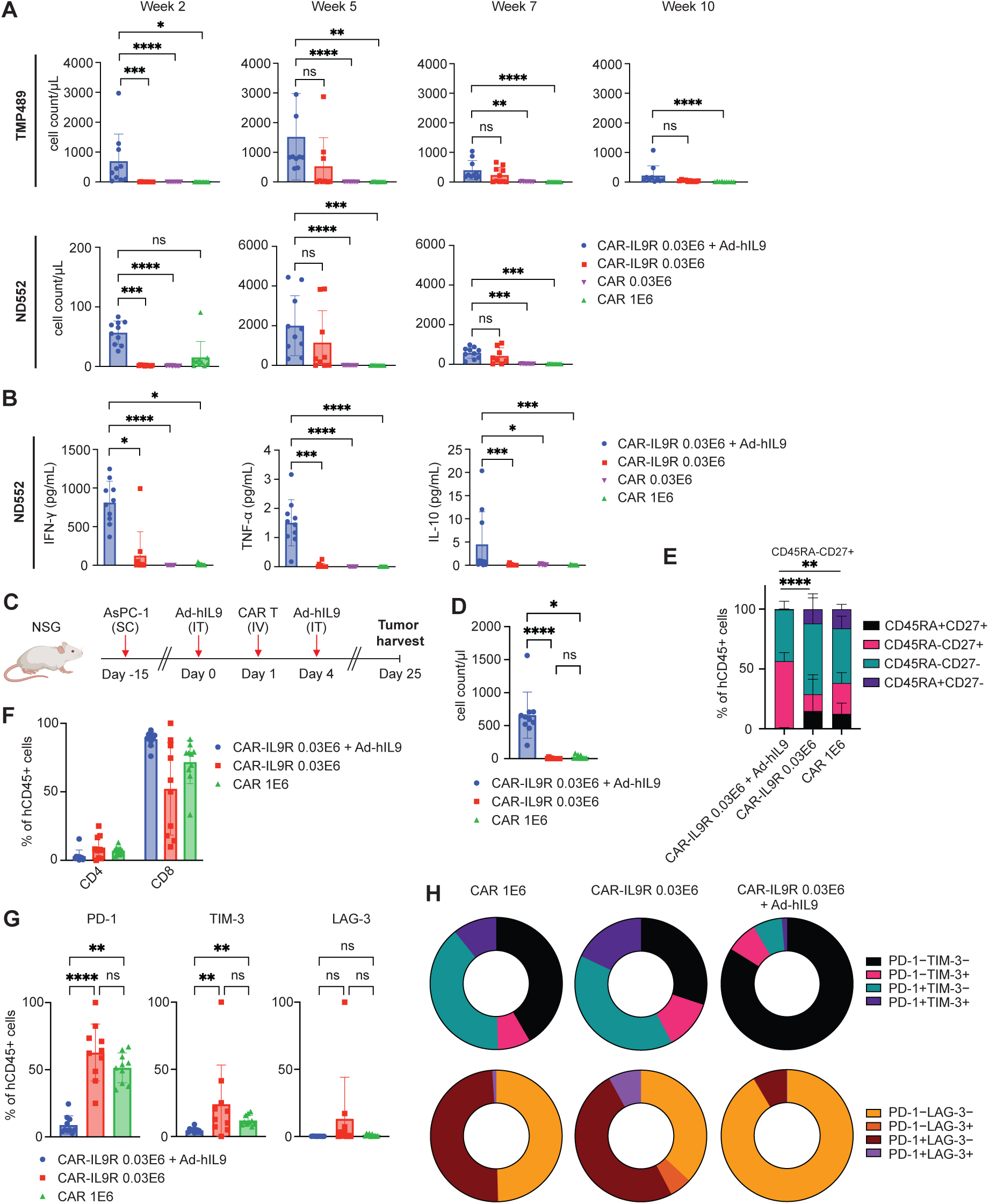
IL-9 signaling enhances CAR T cell expansion, persistence, and tumor infiltration *in vivo*. (A) Enumeration of infused human T cells (hCD45+) in the blood analyzed via flow cytometry (n = 10 per cohort). (B) Concentration of serum cytokines on day 22 after CAR T cell infusion, measured via cytometric bead array (n = 10 per cohort). (C) Schematic of the experimental design used for phenotypic analysis of tumor-infiltrating T cells following treatment. Tumors were collected on day 25 for flow cytometric analysis (Illustration created with BioRender.com). (D) Quantification of infused hCD45+ T cells within tumors, analyzed via flow cytometry (n = 10 per cohort). (E) Flow cytometric analysis of tumor-infiltrating T cell subsets, expressed as percentages of infused hCD45+ cells, identifying CD45RA+CD27+, CD45RA-CD27+, CD45RA-CD27– and CD45RA+CD27-subsets (n= 10 per cohort). (F) Flow cytometry characterization of tumor-infiltrating CD4+ and CD8+ T cells, expressed as percentage of hCD45+ T cells (n = 10 per cohort). (G)Expression levels of co-inhibitory receptors PD-1, TIM-3, and LAG-3 on infused hCD45⁺ T cells within tumors, analyzed via flow cytometry (n= 10 per cohort). (H) Pie charts illustrating the distribution of PD-1, TIM-3, and LAG-3 expression profiles among infused hCD45+ T cells in the tumors, as determined by flow cytometry. The top row displays the proportions of PD-1−TIM-3−, PD-1−TIM-3+, PD-1+TIM-3−, and PD-1+TIM-3+ subsets, while the bottom row shows the proportions of PD-1−LAG-3−, PD-1−LAG-3+, PD-1+LAG-3−, and PD-1+LAG-3+ subsets. Error bars indicate mean ± SD. Kruskal-Wallis one-way ANOVA was used for statistical analysis. ns (not significant) *P* > 0.05; \**P* ≤ 0.05; \*\**P* ≤ 0.01; \*\*\**P* ≤ 0.001; \*\*\*\**P* ≤ 0.0001. See also Figure S3.

To assess the impact of IL-9 signaling on the phenotype of intratumoral CAR T cells, we harvested tumors from our xenograft PDA model (**Figure 3C**) and performed flow cytometry analysis on single-cell suspensions of tumor samples. This analysis revealed a significantly higher infiltration of infused T cells in tumors from mice treated with IL-9-signaling CAR T cells compared to those receiving conventional CAR T cells (**Figure 3D**). Furthermore, tumor-infiltrating T cells from the CAR-IL9R + Ad-hIL9 group exhibited an increased proportion of central memory (CD45RA-CD27+) T cells (T_CM_) (**Figure 3E**) and all treatment groups showed a higher percentage of CD8+ compared to CD4+ T cells (**Figure 3F**).

Tumor-infiltrating T cells from the CAR-IL9R + Ad-hIL9 group had lower expression of the co-inhibitory receptors PD-1 and TIM-3 (**Figure 3G**). Additionally, there was a notable reduction in the percentage of double-positive PD-1+TIM-3+ and PD-1+LAG-3+ T cells, along with a corresponding increase in double-negative PD-1−TIM3− and PD-1−LAG-3− T cells (**Figure 3H**). Blood samples harvested from these mice at the same timepoint also reflected similar trends in CAR T cell phenotype (**Figure S3A-S3E**), reinforcing the findings from the tumors.

### IL-9 signaling enhances CAR T cell function and sustains surface CAR expression, CD4+ T cell levels, and CD8+ T_CM_ phenotype under antigen stress

To explore the mechanisms driving the enhanced antitumor efficacy of IL-9-signaling CAR T cells, we conducted *in vitro* studies on CAR-IL9R T cells. CAR and IL-9R transduction efficiency was confirmed by cell surface staining using flow cytometry (**Figure 4A**). Consistent with results observed in mouse T cells, human CAR-IL9R T cells activated by IL-9 exhibited dose-dependent phosphorylation of STAT1, STAT3, and STAT5 (**Figure 4B**). We next investigated the effects of IL-9 signaling in the context of repeated antigen stimulation, using a continuous antigen exposure (CAE) assay. CAR-IL9R T cells were co-cultured with AsPC-1 cells every other day for 10 days (**Figure 4C**), after which they were harvested for functional assays and flow cytometry phenotyping. CAR-IL9R T cells treated with IL-9 during the CAE assay (IL-9) were compared to cells subjected to this assay in the absence of IL-9 treatment (referred to as Mock-treated cells). The CAR-IL9R T cell baseline product, i.e., cells that had not undergone CAE (D0), was used as a control. A cytotoxicity assay demonstrated that CAR-IL9R T cells undergoing stress testing in the presence of IL-9 demonstrated superior tumor cell killing compared to stress-tested Mock cells, which lost cytotoxic function (**Figures 4D** and **S4A**). These enhanced cytotoxic effects were antigen-specific, as IL-9-treated cells did not eliminate tumor cells lacking mesothelin expression (**Figures 4D** and **S4A**). Additionally, IL-9-treated cells produced higher levels of IFN-γ compared to baseline D0 and Mock cells. This increase was observed both in the absence of antigen stimulation (**Figure 4E** and **S4B**, left panel) and following co-culture with tumor cells (**Figures 4E** and **S4B**, right panel).

**Figure 4:**
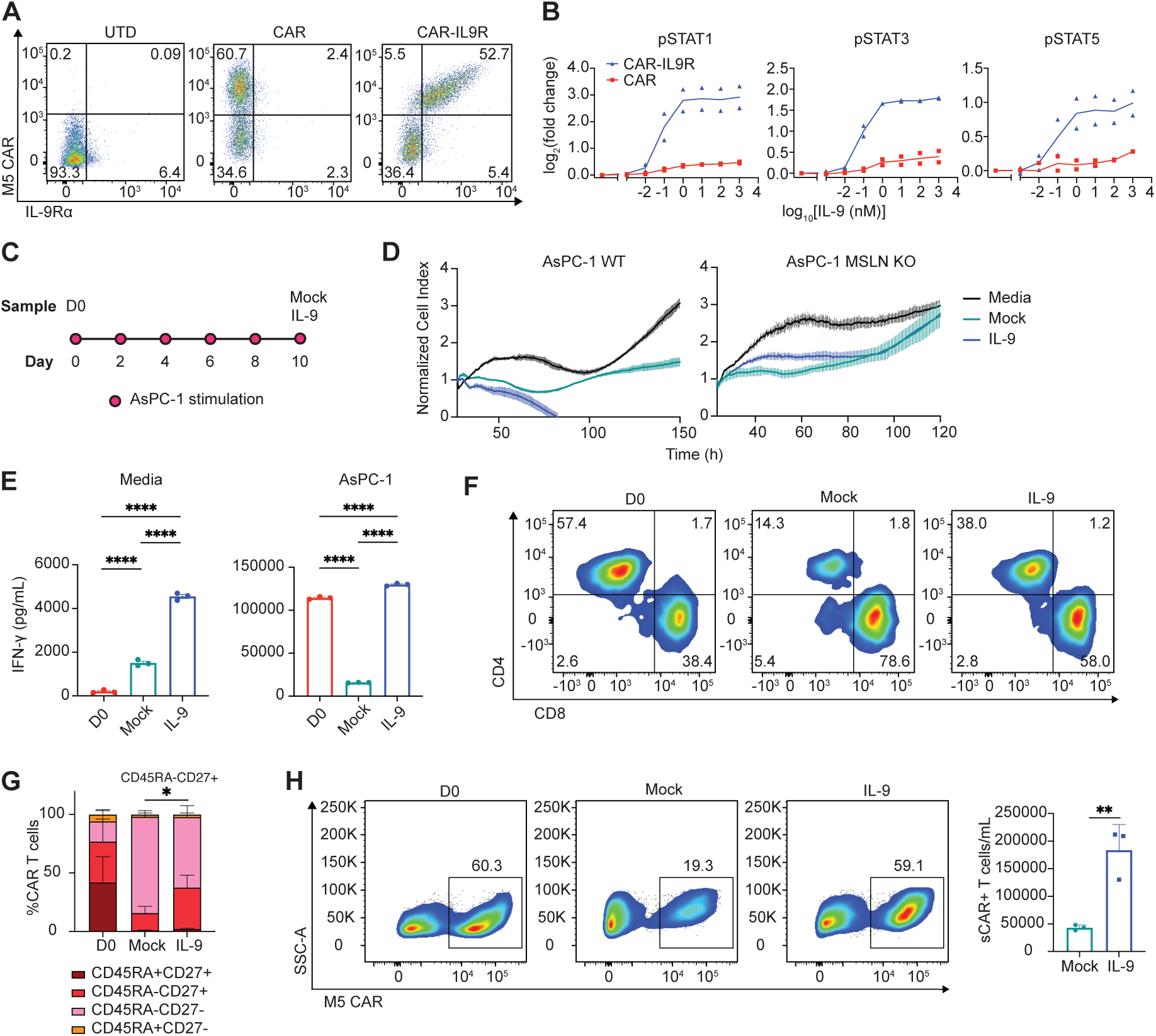
IL-9 signaling enhances CAR T cell function and sustains surface CAR expression, CD4+ T cell levels, and CD8+ T_CM_ phenotype under antigen stress. (A) Flow cytometric analysis of transduction efficiency in human untransduced (UTD), CAR, or CAR-IL9R T cells. (B) Quantification of pSTAT signaling in UTD, CAR, or CAR-IL9R T cells after 20-minute IL-9 stimulation. Data are shown as log fold change of mean fluorescence intensity, with mean and individual values (n = 2 donors). (C) Schematic of the CAR T cell *in vitro* dysfunction model. CAR-IL9R T cells were repeatedly stimulated with AsPC-1 cells every other day for 10 days. Groups include: D0 (initial CAR-IL9R T cell product); Mock (CAR-IL9R cells undergoing the CAE assay in the absence of IL-9); IL-9 (CAR-IL9R cells undergoing the CAE assay in the presence of IL-9). (D) *In vitro* cytotoxicity assay evaluating the lysis of mesothelin (MSLN)-positive wild-type (WT) AsPC-1 tumor cells or MSLN knockout (KO) AsPC-1 cells by IL-9-treated or Mock-treated CAR-IL9R T cells at a 1:4 E:T ratio (n=3). Media was used as a control for tumor growth. Data shown are from normal donor ND552; two additional donors were tested (See Figure S4A). (E) Concentration of interferon-gamma (IFN-γ) produced by D0, Mock or IL-9-treated CAR T cells co-cultured with media (left) or AsPC-1 cells (right), determined by cytometric bead array (n = 3). Data shown are from normal donor ND552; an additional donor was tested (See Figure S4B). (F) Expression of CD4 and CD8 on D0, Mock or IL-9-treated cells. Data shown are from normal donor ND552; two additional donors were tested (See Figure S4C). (G)Flow cytometric analysis of T cell subsets, expressed as percentages of CAR T cells: CD45RA+CD27+, CD45RA-CD27+, CD45RA-CD27-or CD45RA+CD27-(n= 3 donors). (H) Left: surface CAR expression (sCAR+) in D0, Mock or IL-9-treated cells. Data shown are from normal donor ND552; two additional donors were tested (See Figure S4E). Right: quantification of surface CAR expressing T cells per mL following the *in vitro* dysfunction assay (n = 3 donors). Error bars indicate mean ± SD. Significance by two-way ANOVA with Tukey’s post hoc test (E), Fisher’s repeated measures Student’s t test (G) or unpaired t test (H). \**P* ≤ 0.05; \*\**P* ≤ 0.01; \*\*\*\**P* ≤ 0.0001. See also Figures S4 and S5 and Tables S2 and S3.

Flow cytometry analysis showed that, unlike Mock CAR T cells, IL-9-treated CAR T cells maintained balanced CD4+ and CD8+ T cell proportions after repeated antigen stimulation (**Figures 4F** and **S4C**). Additionally, after stress testing, IL-9-treated CAR T cells had an increased percentage of T_CM_ (CD45RA-CD27+) compared with Mock (**Figures 4G** and **S4D**). Notably, while Mock-treated CAR T cells experienced a downregulation of surface CAR expression, which contributes to CAR T cell dysfunction under antigen stress [24, 25], IL-9-treated CAR T cells continued to proliferate and retained surface CAR expression (**Figures 4H** and **S4E**). The continued proliferation and maintenance of CAR expression are recognized as key mechanisms for supporting superior CAR T cell function following prolonged antigen exposure [25].

To further dissect the mechanisms by which IL-9 signaling enhances CAR T cell function under antigen stress, we conducted single-cell RNA sequencing (scRNA-seq). CAR-IL9R T cells from three different donors were subjected to the CAE assay with or without IL-9 treatment (IL-9 and Mock samples, respectively). At day 10, T cells were harvested and isolated via fluorescence activated cell sorting (FACS) of live EpCAM-CD45+ cells, while baseline CAR-IL9R T cells were thawed and sorted for comparison (D0 samples). A total of nine samples were sequenced using the 10X Genomics scRNA-seq platform.

Differential pseudobulk gene analysis of all CAR+ cells revealed that the top upregulated gene in IL-9-treated CAR-IL9R T cells compared to Mock-treated cells was TRERF1, a transcriptional regulatory protein known for its involvement in cholesterol metabolism and steroid hormone signaling [26, 27] (**Figure S5A, Table S2**). Ingenuity pathway analysis (IPA) identified Th1 pathway, IL-20 family signaling, IL-10 signaling, and cholesterol biosynthesis as some of the top pathways enriched in IL-9-treated cells (**Figure S5B, Table S3**). Although IL-10 has traditionally been associated with immunosuppressive roles, recent studies suggest that IL-10 can enhance T cell antitumor immunity by modulating T cell metabolism [28, 29]. Additionally, IL-10 secretion from Th1 cells was shown to be regulated by cholesterol metabolism [30]. Notably, IL-9-signaling CAR T cells exhibited elevated IL-10 secretion both *in vitro* for murine T cells (**Figure 1E**) and *in vivo* for human T cells (**Figure 3B**). This increase in IL-10 production and signaling, coupled with the upregulation of pathways involved in Th1 responses and cholesterol metabolism, prompted us to investigate the potential role of IL-10 in shaping the phenotype and function of IL-9-signaling CAR T cells.

To evaluate whether IL-10 contributes to the antitumor efficacy of IL-9-signaling CAR T cells, we employed a prostate cancer xenograft model. We generated prostate-specific membrane antigen (PSMA)-targeted CAR (Pbbz) T cells co-expressing a dominant-negative TGF-βRII (dnTGF-βRII) and IL-9R (dnTGF-βRII-Pbbz-IL9R) (**Figure S5C**). A clinically tested PSMA dnTGF-βRII CAR (dnTGF-βRII-Pbbz) was used as a control [31]. NSG mice were engrafted with PC3-PSMA tumor cells and received IT Ad-hIL9 along with IV CAR T cell injections. Transduction efficiency was assessed via flow cytometry (**Figure S5D**) and IL-9 production by Ad-hIL9-transduced tumor cells was verified *in vitro* (**Figure S5E**). To evaluate the contribution of IL-10, an additional cohort was treated with an IL-10-neutralizing antibody (α-hIL10) (**Figure S5F**), and IL-10 neutralization was confirmed through serum analysis (**Figure S5G**). Treatment with IL-9-signaling CAR T cells at a 0.03E6 dose significantly reduced tumor progression (**Figure S5H**), improved overall survival (**Figure S5I**), and increased CAR T cell counts in the blood (**Figure S5J**) compared with conventional CAR T cells. Notably, IL-10 neutralization did not diminish antitumor efficacy (**Figure S5H**) or affect blood T cell counts (**Figure S5J**), indicating that IL-10 is not a major driver of the enhanced function of IL-9-signaling CAR T cells.

### IL-9 treatment enhances CAR T cell effector phenotype and reduces dysfunction under antigen stress

To further investigate how IL-9 signaling contributes to the improved antitumor activity of CAR T cells, we conducted detailed analysis of scRNA-seq data. Semi-unsupervised clustering using uniform manifold approximation and projection (UMAP), identified 15 distinct T cell clusters (**Figure 5A, Figure S6**). Analysis of cluster distribution revealed a higher proportion of cells in effector clusters within the IL-9-signaling group (**Figure 5B**). Specifically, CAR+ T cells in the IL-9-treated group showed a significant increase in the CD4+ Activated and *GNLY*++ Effector clusters. In contrast, the Mock group exhibited higher proportions of CAR+ cells in CD8+ Exhausted and *GZMA*++ Cytotoxic clusters (**Figure 5C**). The Day 0 samples displayed typical distributions of naïve and effector cells, as expected from cells that had not yet encountered antigen (**Figures 5B** and **5C**).

**Figure 5:**
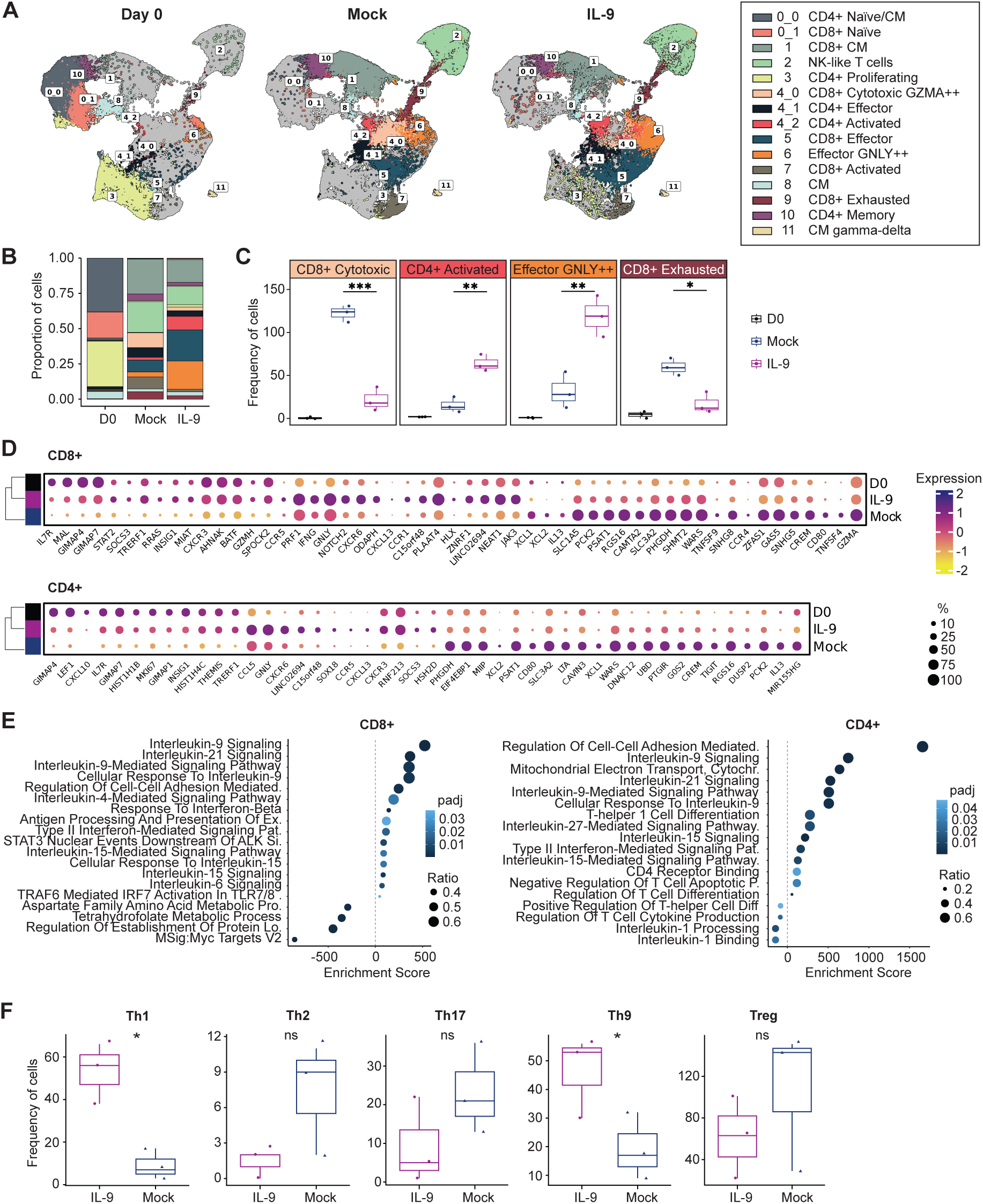
IL-9 treatment enhances CAR T cell effector phenotype and reduces dysfunction under antigen stress. (A) Uniform Manifold and Approximation Projection (UMAP) visualization of scRNA-seq data from Day 0 (D0), IL-9-treated and Mock-treated T cells from donors TMP489, ND552 and ND541. UMAP clusters are color-coded, with annotations shown in the right panel. (B) Bar graph illustrating the proportion of D0, IL-9-treated and Mock-treated T cells by cluster. Data were downsampled uniformly across treatments and clusters to ensure comparability. (C) Boxplots displaying the frequency of CAR+ cells among D0, IL9-treated and Mock-treated groups. Only clusters for which there is a statistically significant difference in the frequency of CAR+ cells between IL9-treated and Mock-treated groups are shown. (D) Dot plot illustrating differentially expressed (DE) genes between D0, IL9-treated and Mock-treated groups within CD4+ and CD8+ CAR+ T cells with adjusted p-values < 0.05. Dot size represents the percentage of cells expressing each gene, while color intensity denotes expression levels. (E) Gene set enrichment analysis of CD4+ and CD8+ CAR+ cells. Dot size reflects the ratio of genes in each pathway, and color intensity denotes statistical significance (padj: adjusted p-value). (F) Frequency distribution of IL9-treated and Mock-treated CD4+ CAR+ subsets, assessed by signature scoring. Cluster composition analysis was performed using Welch’s t-test. DE statistical analysis was performed using MAST. Signature scores were calculated as area under the curve. Donor comparisons between treatment groups were conducted using Fisher’s repeated measures Student’s t-test. ns (not significant) *P* > 0.05; \**P* ≤ 0.05; \*\**P* ≤ 0.01; \*\*\**P* ≤ 0.001; \*\*\*\**P* ≤ 0.0001. See also Figure S6 and Tables S4 and S5.

Next, we investigated differential gene expression in the CD8+ and CD4+ CAR+ compartments (**Figure 5D, Table S4**). Notably, IL-9 treatment in CD8+ T cells led to higher expression of transcription factors such as *TRERF1*, BATF, STAT2, and HLX, along with upregulation of chemokine-related genes like *CXCL13*, *CCR1*, *CCR5*, *CXCR6*, and *CXCR3*. Markers of naïve or memory CD8+ T cells, such as *IL7R*, and cytotoxicity-related genes like *GNLY*, *IFNG*, *GZMH*, and *PRF1*,were also upregulated. In contrast, expression of genes linked to activation, including *GZMA*, *TNFSF4* (OX-40L), *CD80*, and *TNFSF9* (4-1BBL), was reduced. Since sustained expression of T cell activation genes is often linked to exhaustion states [32], their downregulation in IL-9-treated cells may indicate decreased dysfunction. In CD4+ T cells, IL-9 treatment similarly upregulated chemokine-related genes including *CXCL10*, *CCL5*, *CXCR6*, *CCR5*, *CXCL13*, and *CXCR3*, along with markers of naïve/memory T cells (*LEF1*, *IL7R*). Additionally, there was increased expression of proliferation-associated genes such as *MKI67* and cytotoxicity-related genes like *GNLY*, while *CD80* and *TIGIT* were downregulated. Unsupervised hierarchical clustering revealed that IL-9-treated CAR+ cells retained a transcriptional profile more closely aligned with Day 0 samples than the Mock group for both CD8+ and CD4+ CAR+ cells (**Figure 5D**), suggesting that IL-9 treatment promotes less differentiated T cell states compared to Mock.

Gene set enrichment analysis (GSEA) revealed significant enrichment of interleukin signaling pathways in IL-9-treated CD8+ CAR+ T cells, including IL-4, IL-6, IL-15, IL-21, and type II interferon responses. A similar pattern was observed in CD4+ CAR+ cells, where IL-21, IL-27, IL-15 and type II interferon signaling pathways were significantly enriched, along with those promoting Th1 differentiation and cell-cell adhesion (**Figure 5E, Table S5**). Signature scoring confirmed an elevated frequency of Th1 cells and Th9 cells in the IL-9-treated group, with no notable changes in Th2, Th17, or Treg populations (**Figure 5F**).

### IL-9 signaling directs CAR+ CD8+ T cells towards central memory clusters and away from dysfunctional clusters under antigen stress

We investigated the impact of IL-9 on CAR T cell differentiation by analyzing CAR+ cell trajectories along pseudotimes using a semi-unsupervised approach [33]. We identified six distinct trajectories: three for CD8+ cells, two for CD4+ cells, and one combined CD8+/CD4+ (**Figures 6** and **S7A**), with four trajectories showing significant differences between IL-9-treated and Mock conditions (**Figure 6A**).

**Figure 6:**
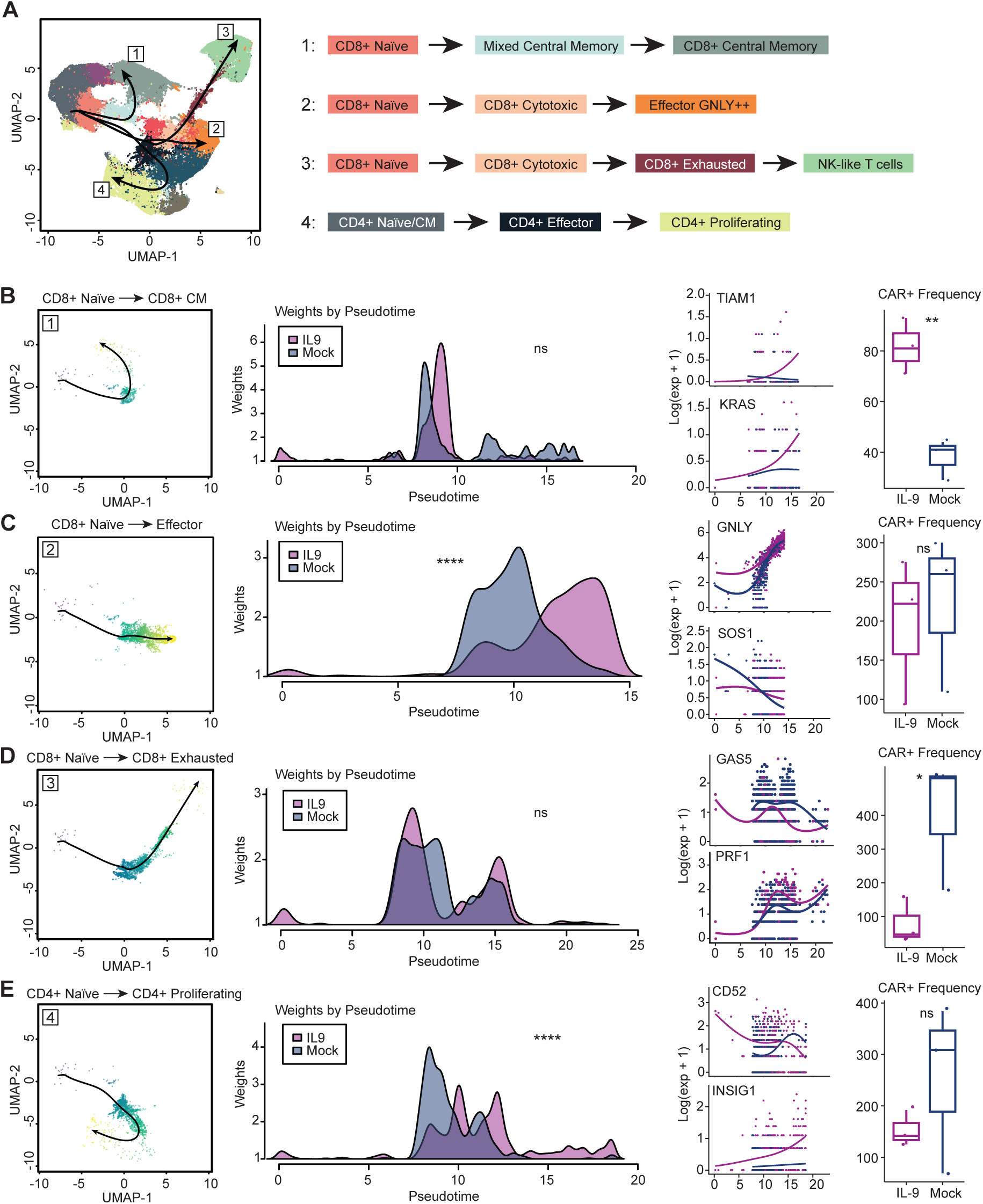
IL-9 signaling directs CAR+ CD8+ T cells towards central memory clusters and away from dysfunctional clusters under antigen stress. (A) Uniform Manifold and Approximation Projection (UMAP) visualization of four identified trajectories overlayed on clusters, with corresponding cluster paths diagramed on the right. (B) CD8+ Naïve to CD8+ Central Memory (CM) trajectory: Left: UMAP; Middle: weights by pseudotime; Right: differential expression of select genes along the trajectory, and CAR+ cell distribution frequency along the trajectory. (C) CD8+ Naïve to CD8+ Effector GNLY++ trajectory: Left: UMAP; Middle: weights by pseudotime; Right: differential expression of select genes along the trajectory, and CAR+ cell distribution frequency along the trajectory. (D) CD8+ Naïve to CD8+ Exhausted trajectory: Left: UMAP; Middle: weights by pseudotime; Right: differential expression of select genes along the trajectory, and CAR+ cell distribution frequency along the trajectory. (E) CD4+ Naïve to CD4+ Proliferating trajectory: Left: UMAP; Middle: weights by pseudotime; Right: differential expression of select genes along the trajectory, and CAR+ cell distribution frequency along the trajectory. Weighted pseudotime comparisons were performed by Wilcoxon rank-sum test with continuity correction. Differential gene expression along trajectories were calculated using a negative binomial generalized additive model (NB-GAM). Comparisons of CAR+ cell frequencies between treatments were conducted by Fisher’s repeated measures Student’s t-test. ns (not significant) *P* > 0.05; \**P* ≤ 0.05; \*\**P* ≤ 0.01; \*\*\**P* ≤ 0.001; \*\*\*\**P* ≤ 0.0001. See also Figure S7.

In the first CD8+ trajectory, cells progressed from naïve to central memory populations (**Figure 6A** and **6B**). IL-9 treatment increased the number of CAR+ cells along this path, without altering the distribution of cell weights along the trajectory. This data aligns with the observed increase in the proportion of tumor-infiltrating T_CM_ in mice from the xenograft PDA model treated with IL-9-signaling compared to conventional CAR T cells (**Figure 3E**). Differential expression analysis identified genes statistically different along the trajectory with the addition of IL-9, including genes involved in migration (*TIAM1*) and effector function (*KRAS*) that increased over pseudotime compared to Mock (**Figure 6B, Figure S7B**). In the second CD8+ trajectory, cells progressed from a naïve state through cytotoxic stages, culminating in T cells with robust effector function (**Figure 6A** and **6C**). IL-9 treatment influenced cell distribution along the pseudotime, resulting in more IL-9-signaling CAR T cells reaching a strong effector state, consistent with the increased proportion of cells in the Effector *GNLY*++ cluster (**Figure 5C**). In this trajectory, IL-9 treatment led to increased expression of cytotoxicity related genes (*GNLY*, *GZMB*), whereas low expression of genes involved in T cell differentiation (*SOS1*) and regulation of T cell responses (*TNFSF13B*) was maintained along the pseudotime (**Figure 6C, Figure S7B)**. The third CD8+ trajectory showed cells progressing from naïve through cytotoxic states to exhausted and dysfunctional states resembling NK-like cells (**Figure 6A** and **6D**). This progression, consistent with findings in our solid tumor model of CAR T cell dysfunction, occurs when continuous antigen exposure drives CD8+ T cells to transition into NK-like states [24]. Notably, fewer cells followed this path in the IL-9-treated group, suggesting that IL-9 may redirect CAR T cell progression away from an exhausted state. This aligns with the reduced proportion of IL-9-signaling T cells observed in the CD8+ Exhausted cluster (**Figure 5C**). Multiple genes differ by treatment, including *GAS5, PRF1,* and chemokine/cytokine related genes (*CCL5, IL13, CCL1*) (**Figure 6D**, **Figure S7B**).

The CD4+ trajectory, trajectory 4, transitioned from naïve to effector and ultimately to proliferating states (**Figure 6A** and **6E)**. IL-9 treatment promoted this cycling, as more cells reached the trajectory’s terminal point. Conversely, Mock-treated CAR+ cells stalled in effector states. This suggests that IL-9 actively supports a return to proliferative states in CD4+ CAR T cells. Differentially expressed genes along this trajectory included decreased expression of *CD52*, involved in regulating T cell responses, and increased expression of the cholesterol biosynthesis gene *INSIG1* and proliferation marker *MKI67* (**Figure 6E**, **Figure S7B**).

Together, these findings indicate that IL-9 signaling influences CAR T cell differentiation under antigen stress, enhancing CD8+ CAR T cell progression toward central memory and effector states, and promoting proliferation in CD4+ cells.

### RNA velocity and transcription factor analyses provide mechanistic insight into how IL-9 signaling redirects CAR T cell fate

To independently validate our trajectory findings, we performed RNA velocity analysis to study changes in gene expression over time. RNA velocity defines vectors that predict the future state of individual cells based on the ratio of spliced and unspliced mRNA, with the assumption that more differentiated cells have a higher proportion of spliced mRNA [34]. We found that spliced mRNA progressively increased from Day 0 (40%) to IL-9-treated (42%) and Mock-treated cells (44%), indicating a higher proportion of differentiated cells in the Mock-treated group (**Figure 7A**).

**Figure 7:**
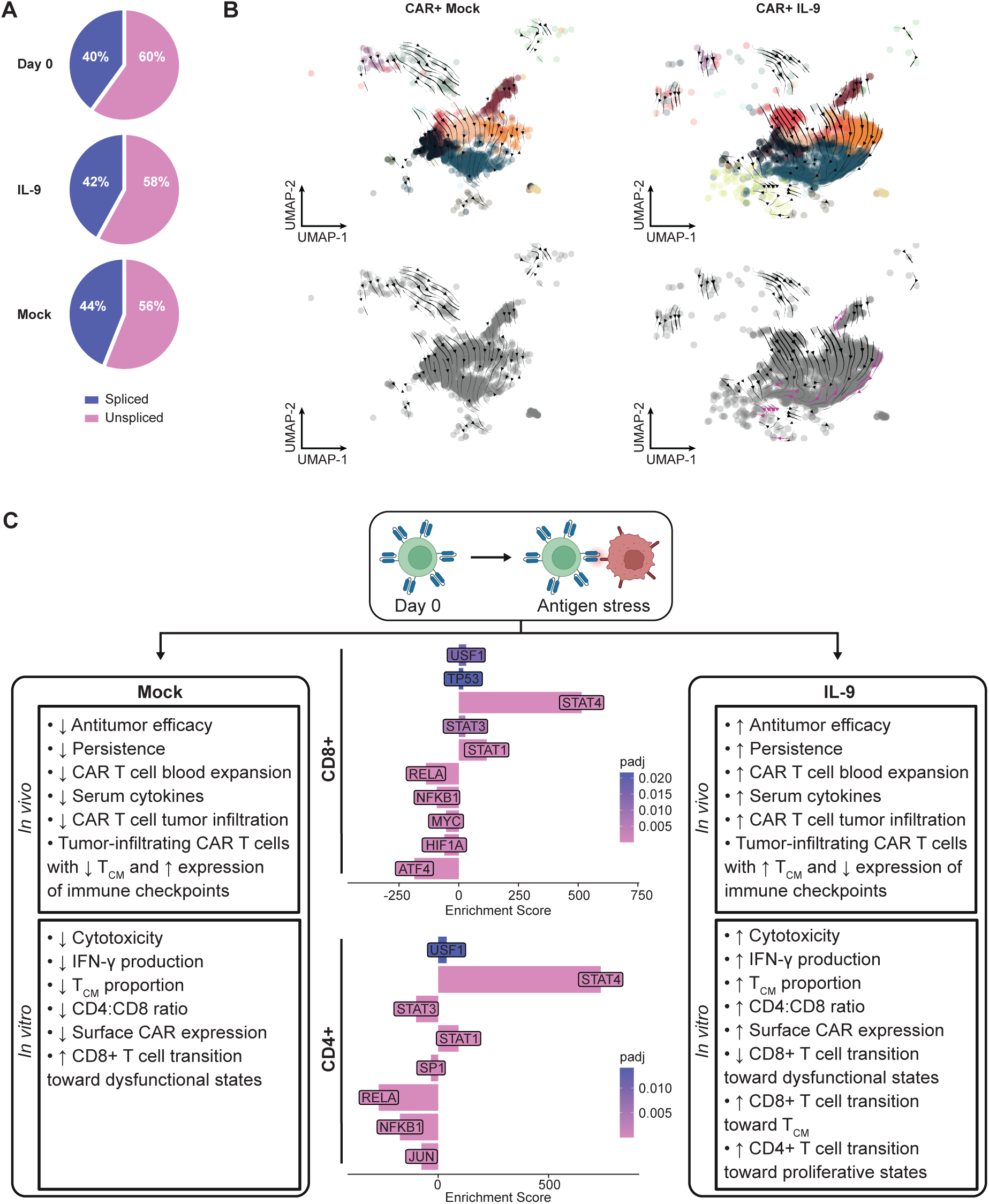
RNA velocity and transcription factor analyses provide mechanistic insights into how IL-9 signaling redirects CAR T cell fate. (A) Pie charts illustrating the fractions of spliced and unspliced reads in T cells for D0, IL-9-treated and Mock-treated groups. (B) Velocity fields projected onto UMAPs for IL-9-treated and Mock-treated CAR+ cells. The top panels display UMAPs colored by clusters, while the bottom panels show black-and-white UMAPs to enhance the visualization of velocity vectors. Violet arrows indicate vectors predicting future states that shift from right-to-left and/or upward. (C) Schematic overview comparing the phenotype of CAR T cells under antigen stress in the absence (Mock, left panel) or presence of IL-9 signaling (IL-9, right panel), alongside the transcription factors potentially driving this phenotype (middle panel). T_CM_, central memory T cells. Differentially expressed genes with a log fold change greater than ±1 were analyzed using the TRRUST database, with results filtered to include only those with an adjusted p-value (padj) ≤ 0.02. TRRUST (Transcriptional Regulatory Relationships Unraveled by Sentence-based Text-mining) is a manually curated database of human and mouse transcriptional regulatory networks, providing information on transcription factor-target interactions and modes of regulation. See also Figure S7 and Tables S6 and S7.

Further RNA velocity analyses reveal that IL-9 signaling significantly altered the rate and direction of change of CAR T cells exposed to antigen stress over time. In Mock-treated CAR T cells, velocity vectors predominantly followed a path from the top left to the bottom right of the UMAP plot (**Figure 7B**). In contrast, in IL-9-treated CAR T cells, velocity vectors also showed a distinct horizontal progression, marked by violet-colored vectors, indicating movement away from terminal states and toward less differentiated fates. This horizontal directional progression includes vectors laterally exiting the Exhaustion cluster and vectors progressing from an Effector *GNLY*++ cluster toward a predicted future CD4+ proliferating state. Overall, RNA velocity predicts that IL-9 signaling drives CAR T cells toward less differentiated and more proliferative effector states under antigen stress compared to conventional CAR T cells (**Figure 7B**).

Both trajectory and RNA velocity analyses indicate that IL-9 signaling promotes the transition of CAR T cells toward functional rather than dysfunctional states. This transition fosters the development of proliferative and effector T cells under antigen stress. Despite relying on distinct inputs, the results from these two methods are consistent: RNA velocity employs the relative abundance of spliced versus unspliced mRNA to infer the rate and direction of cell-state transitions, whereas trajectory analysis relies on a count matrix and minimum spanning tree (MST) modeling. Together, these complementary approaches validate our conclusion that IL-9 signaling promotes CAR T cell memory and proliferation over terminal differentiation, promoting the dramatically enhanced potency of CAR-IL9R T cells.

To further investigate the transcriptional mechanisms underlying the phenotypic differences between IL-9-treated and Mock-treated CAR T cells, we analyzed regulatory interactions between transcription factors (TFs) and target genes from our scRNA-seq data using the TRRUST human database (**Table S6**) [35]. Results were confirmed using IPA upstream regulatory analysis (**Table S7**). These analyses identified transcriptional regulatory networks (**Figure 7C**, middle panel) that may explain the differences observed in Mock-treated (**Figure 7C**, left panel) and IL-9-treated (**Figure 7C**, right panel) CAR-IL9R T cells under antigen stress. IL-9 treatment strongly enriched STAT4 activity in both CD4+ and CD8+ CAR T cells (**Figures 7C** and **S7C**). This was unexpected because STAT4 has not previously been associated with IL-9 signaling. Additionally, STAT4 activity was elevated in IL-9-treated and baseline CAR T cells compared to Mock, specifically within cluster 9 (CD8+ Exhausted) (**Figures S7C** and **S7D**). Both STAT1 and STAT3 are established mediators of γ_c_ cytokine signaling, including IL-9, while STAT4 traditionally supports Th1 responses via IL-12 signaling. IL-9 treatment also enriched STAT1 activity, known to mediate IFN-γ functions [36]. The USF1 TF pathway was also upregulated in both CD8+ and CD4+ CAR T cells, however the STAT3 pathway was only upregulated in CD8+ CAR T cells. STAT3 signaling has been shown to drive T cell memory formation [37]. Additionally, the USF1 TF, which regulates genes involved in sugar and lipid metabolism [38, 39] may provide a proliferative advantage to IL-9-signaling CAR T cells.

Both CD8+ and CD4+ IL-9-signaling CAR T cells exhibit downregulation of TF pathways associated with T cell activation (RELA, NFkB1) under antigen stress, potentially reducing T cell dysfunction. Further, IL-9-signaling CD8+ CAR T cells exhibit decreased levels of ATF4, HIF-1α and MYC pathways, suggesting a lower cellular stress response. These results provide mechanistic insight into how IL-9 signaling reprograms CAR T cell differentiation upon repeated antigen exposure, enhancing central memory formation and reducing exhaustion in CD8+ T cells while fostering a proliferative effector phenotype in CD4+ T cells to potentiate antitumor efficacy, CAR T cell expansion and persistence *in vivo* (summarized in **Figure 7C)**.

In summary, IL-9 signaling redirects CD4+ and CD8+ CAR T cell fate, primarily through STAT1 and STAT4 signaling to enable less differentiated and exhaustion-resistant states, enhancing their efficacy against solid tumors.

## DISCUSSION

A major obstacle hindering the effectiveness of CAR T cell therapies targeting solid tumors, which comprise over 90% of diagnosed cancers [40], is the immunosuppressive TME. CAR T cells exhibit limited tumor infiltration into the TME, and those that do penetrate the tumor often become dysfunctional [41]. High doses of CAR T cells have been associated with on-target toxicities [42, 43], and this study addresses these critical barriers by endowing CAR T cells with IL-9 signaling, which is typically limited in T cells (Jiang et al., 2024), to enhance antitumor efficacy. IL-9-signaling CAR T cells exhibit superior expansion, persistence, and tumor infiltration, resulting in enhanced antitumor efficacy even at low CAR T cell doses. IL-9 signaling in the context of CAR T cells has the unusual ability to induce multiple differentiation trajectories including memory, effector, and proliferative states. This approach potentially reduces the risks associated with overactivation and minimizes toxicities in off-tumor tissues, offering a promising strategy for enhancing clinical outcomes in solid tumors.

Our findings reveal that IL-9 signaling uniquely influences the fate of CD4+ CAR T cells. Transcriptomic analysis shows an enrichment of activated CD4+ T cell clusters. Trajectory and RNA velocity analyses indicate that IL-9-treated CAR T cells progress toward less differentiated, proliferative CD4+ clusters, suggesting enhanced proliferative potential in response to IL-9 signaling. Notably, IL-9-signaling CAR T cells preserved a balanced CD4^+^ to CD8^+^ T cell ratio, which may enhance their functional resilience, as CD4^+^ T cells are crucial for sustaining T cell responses under chronic antigen exposure [44]. Higher CD4^+^ to CD8^+^ T cell ratios have been linked with increased IFN-γ production following CAR T cell transfer, correlating with more favorable clinical outcomes [45]. Additionally, CD4^+^ T cells contribute to the durability of CAR T cell responses, supporting long-term efficacy [46, 47].

IL-9 signaling promotes CD4+ CAR T cell differentiation toward Th1 cells, a process accompanied by increased secretion of IFN-γ and TNF-α, both critical for antitumor responses. The transcription factor STAT4 is essential for Th1 differentiation and the production of pro-inflammatory cytokines; STAT4-deficient mice exhibit a profound Th1 defect [48, 49]. Our data suggest that in IL-9-signaling CAR T cells, IL-9, rather than IL-12, promotes Th1 differentiation through induction of the STAT4 pathway. These findings align with a recent study in acute myeloid leukemia, where IL-9 secretion by leukemia stem cells induces Th1 polarization [50]. Th1 cells and their cytokines are strongly associated with better clinical outcomes across multiple cancer types [51], and Th1 differentiation is linked to improved efficacy in adoptive cell therapies [52, 53]. Notably, IL-9-signaling CAR T cells also retain a population of Th2-polarized CAR T cells. Recent work indicates that elevated Th2 functionality in CAR T infusion products correlates with long-term CAR T cell persistence and superior outcomes in pediatric hematologic malignancies [54]. In our study, IL-9 treatment also increases the percentage of Th9 cells, which have been implicated in antitumor responses in preclinical solid tumor models [16, 17]. Collectively, CD4+ IL-9-signaling CAR T cells, polarized toward Th1 and Th9 cells while maintaining Th2 cells, exhibit superior antitumor efficacy and persistence both *in vitro* and *in vivo*.

Our findings show that IL-9 signaling directs CD8+ CAR T cells toward less differentiated T cell fates. Tumor-infiltrating IL-9-signaling CAR T cells display an enhanced central memory phenotype, along with increased cytokine secretion and reduced dysfunction compared to conventional CAR T cells. Transcriptomic analyses reveal that IL-9 signaling drives CD8+ T cells toward central memory clusters, while reducing the representation of dysfunctional CD8+ clusters. Further, velocity vectors of IL-9-treated CAR T cells show distinct horizontal progression, indicating movement away from terminal dysfunctional states toward less differentiated fates, unlike the vectors of conventional CAR T cells. Importantly, the enrichment of memory-like features has been strongly associated with long-term remission in CAR T cell patients [55, 56], whereas CAR T cell dysfunction negatively correlates with clinical efficacy [57]. Consistent with this, we observed that IL-9-signaling CAR T cells had potent effects in our xenograft PDA and prostate tumor models. The CAR T cells persisted in circulation for up to ten weeks and tumor rechallenge led to rapid tumor rejection, underscoring the potential of IL-9 signaling to enhance the function and persistence of CAR T cells.

The role of STAT transcription factor pathways in mediating increased central memory formation and decreased dysfunction under antigen stress in CD8+ IL-9-signaling CAR T cells remains to be fully elucidated. Analysis of differentially expressed genes in CAR T cells after stress testing by TRRUST and IPA reveals a significant upregulation of the STAT4 transcription factor pathway following IL-9 treatment. Comparison of STAT4 activity score from baseline, Mock and IL-9 groups illustrates that IL-9-treated and baseline CAR T cells exhibit higher levels of STAT4 activity in cluster 9 (CD8+ Exhausted) than Mock, suggesting that STAT4 may influence the frequency of IL-9-signaling CAR T cells in the exhaustion cluster. STAT1 and STAT3 TF pathways are also upregulated in IL-9-signaling CD8+ CAR T cells after continuous antigen exposure. It is reasonable to hypothesize that the relative activation of these pathways likely influences CAR T cell fate. A recent study provides evidence that STAT1 functions as a rheostat to control the balance of T cells in a stem-like state with proliferative capacity and a terminally differentiated effector state (Jiang et al., 2024). STAT3 signaling plays a role in the development of memory T cells [37]. Although the STAT5 TF pathway has also been shown to be activated upon IL-9-signaling, we did not find evidence of high levels of activation after antigen stress. It is possible that Mock-treated cells produce sufficient IL-2, explaining the similarity of STAT5 activation in control and IL-9 treated groups.

In summary, we engineered CAR T cells to incorporate IL-9 signaling and assessed the impact on antitumor efficacy and phenotypic characteristics. Notably, even at low doses, IL-9 signaling significantly enhanced CAR T cell expansion, persistence, and tumor infiltration, leading to improved tumor clearance and prolonged survival in *in vivo* models of pancreatic and prostate cancer. Under antigen stress, IL-9 fosters a favorable phenotype, characterized by enhanced central memory formation, reduced expression of ATF4 and reduced dysfunction in CD8+ CAR T cells. Coupled with increased effector proliferation in CD4+ CAR T cells, this yields less differentiated and more potent CAR T cells. These findings position IL-9 signaling as a promising strategy to overcome key barriers in CAR and TCR T cell therapies targeting solid tumors.

## ACKNOWLEDGEMENTS

We would like to thank Donna Gonzales, Tong Da, Carolyn Shaw, Yujie Ma, John Scholler, Ting-Jia Fan, Andrew Rech and Enrico Radaelli for technical assistance and/or helpful discussions, and John Tobias for technical assistance with Ingenuity Pathway Analysis (IPA). We also thank the Translational and Correlative Studies Laboratory (TCSL) at the University of Pennsylvania for their generous provision of patient serum samples. We acknowledge the Human Immunology Core (HIC) at University of Pennsylvania for reliable supply of healthy human T cells and serum and for performing the Luminex cytokine assays, the Genomics Facility at the Wistar Institute for RNA extraction and scRNA-seq assays, and the Comparative Pathology Core (CPC) at School of Veterinary Medicine for pathological assessment, immunohistochemistry, image acquisition, and analysis. The veterinary pathologist performing the histopathological analysis is part of the University of Pennsylvania Penn Vet Comparative Pathology Core Facility (RRID:SCR_022438) and is supported by the Abramson Cancer Center Support Grant (P30 CA016520). The scanner used for whole slide imaging and the image analysis software was supported by an NIH Shared Instrumentation Grant (S10 OD023465-01A1). The Leica BOND RXm instrument used for IHC was acquired through the Penn Vet IIZD Core pilot grant opportunity 2022. Funding was provided by the Mark Foundation (19-011MIA), Prostate Cancer Foundation (PCF) TACTICAL grant (22-TACT03), and Parker Institute for Cancer Immunotherapy (PICI) (C-04339). Additionally, S.C. is supported by the PICI RISE Scholar Award. U.U. is supported by a Mildred-Scheel-Postdoctoral Fellowship of the German Cancer Aid.

## AUTHOR CONTRIBUTIONS

Conceptualization: S.C., R.M.Y., C.H.J., Methodology: S.C., R.M.Y., W.V.W., Investigation: S.C., U.U., A.V.F., C.A.A., S.A., M.S., Formal analysis: S.C., W.V.W., Visualization: S.C., W.V.W., Funding acquisition: S.C., R.M.Y., C.H.J., Project administration: S.C., R.M.Y., Supervision: R.M.Y., C.H.J., Writing – original draft: S.C., Writing – review & editing: S.C., R.M.Y., C.H.J., Writing – editing: W.V.W., U.U., A.V.F., C.A.A., S.A., M.S.

## FIGURE LEGENDS: SUPPLEMENTARY FIGURES

**Figure S1: IL-9 signaling enhances CAR T cell infiltration into tumor and spleen without markedly altering the TME, related to Figure 1**.

(A) Representative RNA in situ hybridization photomicrographs (40x magnification; scale bar: 40 μm) indicating the presence of infused T cells and mesothelin (MSLN)-expressing tumor cells.

(B) Quantification of infused CD45.1+ T cells in spleen single-cell suspensions (left) or blood samples (right), analyzed via flow cytometry (n = 5 per cohort).

(C) Serum cytokine concentrations on days 5 and 6 post-treatment, analyzed via Luminex assay (n = 10 per cohort).

(D) Enumeration of various cell populations within tumor single-cell suspensions, as indicated, analyzed via flow cytometry (n = 5 per cohort). Tumors were harvested on day 6 post-treatment.

(E) Quantification of specified cell populations within tumor sections, analyzed via immunohistochemistry (n = 5 per cohort). Tumors were harvested on day 5 post-treatment.

Error bars indicate mean ± SD. Statistical analysis was performed using Kruskal-Wallis one-way analysis of variance. ns (not significant) *P* > 0.05; \**P* ≤ 0.05.

**Figure S2: Murine IL-9 activates IL-9R in human CAR-IL9R T cells without exogenous addition of IL-9, related to Figure 2**.

(A) Concentration of human IL-9 (hIL-9, left) or murine IL-9 (mIL-9, right) in serum collected on day 22 after CAR T cell infusion, analyzed via cytometric bead array (n = 10 per cohort).

(B) Schematic of experimental design (created with BioRender.com). SC, subcutaneous; IT, intratumoral; IV, intravenous; IP, intraperitoneal; α-mIL9, neutralizing antibody targeting mIL-9.

(C) Kaplan-Meier survival curves. For donor TMP489: n = 10 for Ad-hIL9 and CAR-IL9R 0.03E6 + α-mIL9; n = 7 for CAR-IL9R 0.03E6 + isotype control; n = 9 for CAR-IL9R 0.03E6 + Ad-hIL9. For donor ND552: n = 10 per cohort.

(D) Tumor volume progression over time.

(E) Count of infused T cells (hCD45+) in the blood, analyzed via flow cytometry. For Ad-hIL9 and CAR-IL9R 0.03E6 + α-mIL9: n = 10; for CAR-IL9R 0.03E6 + isotype control: n = 7; for CAR-IL9R 0.03E6 + Ad-hIL9: n = 9.

Error bars indicate mean ± SD. Statistical significance was calculated using the log-rank Mantel-Cox test or Kruskal-Wallis one-way ANOVA. ns (not significant) *P* > 0.05; \**P* ≤ 0.05; \*\**P* ≤ 0.01; \*\*\**P* ≤ 0.001; \*\*\*\**P* ≤ 0.0001.

**Figure S3: IL-9-signaling induces a central memory phenotype and reduces co-inhibitory receptor expression in CAR T cells, related to Figure 3**.

(A) Enumeration of infused human T cells (hCD45+) in the blood harvested on day 25 post-CAR T cell infusion, analyzed via flow cytometry (n = 10 per cohort).

(B) Flow cytometric analysis of T cell subsets within the infused hCD45+ population in the blood, categorized as CD45RA+CD27+, CD45RA-CD27+, CD45RA-CD27– and CD45RA+CD27-(n= 10 per cohort).

(C) Proportions of CD4+ and CD8+ T cells expressed as percentage of infused T cells (hCD45+) in the blood, determined by flow cytometry (n= 10 per cohort).

(D) Expression levels of co-inhibitory receptors PD-1, TIM-3 and LAG-3 on infused hCD45+ T cells in the blood, expressed as percentage of infused T cells (n= 10 per cohort.

(E) Pie charts illustrating the distribution of PD-1, TIM-3, and LAG-3 expression profiles among infused hCD45+ T cells in the blood, as determined by flow cytometry. The top row displays the proportions of PD-1−TIM-3−, PD-1−TIM-3+, PD-1+TIM-3−, and PD-1+TIM-3+ subsets, while the bottom row shows the proportions of PD-1−LAG-3−, PD-1−LAG-3+, PD-1+LAG-3−, and PD-1+LAG-3+ subsets.

Error bars indicate mean ± SD. Statistical significance was calculated using Kruskal-Wallis one-way ANOVA. ns (not significant) *P* > 0.05; \**P* ≤ 0.05; \*\**P* ≤ 0.01; \*\*\**P* ≤ 0.001.

**Figure S4: IL-9 signaling enhances CAR T cell effector function and promotes a central memory phenotype after repeated antigen stimulation across multiple donors, related to Figure 4**.

(A) *In vitro* cytotoxicity assay evaluating the lysis of mesothelin (MSLN)-positive wild-type (WT) AsPC-1 tumor cells or MSLN knockout (KO) AsPC-1 cells by IL-9-treated or Mock-treated CAR-IL9R T cells at a 1:4 E:T ratio (n=3). Media was used as a control for tumor growth.

(B) Concentration of interferon-gamma (IFN-γ) secreted by CAR T cells at baseline (D0), and following Mock or IL-9 treatment, after co-culture with media (left) or AsPC-1 cells (right), determined by cytometric bead array (n = 3).

(C) Flow cytometric analysis of CD4 and CD8 expression levels in CAR T cells at baseline (D0), and after Mock or IL-9 treatment.

(D) Assessment of CD45RA and CD27 expression on CAR T cells at baseline (D0) and following Mock or IL-9 treatment.

(E) Evaluation of surface CAR expression on T cells at baseline (D0), and after Mock or IL-9-treatment.

Error bars indicate mean ± SD. Statistical significance was determined by two-way ANOVA with Tukey’s post hoc test.

**Figure S5: IL-9-driven enhancement of CAR T cell antitumor efficacy in a prostate cancer xenograft model is independent of IL-10 signaling, related to Figure 4**.

(A) Volcano plot identifying the most significant differentially expressed genes between IL-9-treated and Mock-treated CAR T cells. Red dots represent genes with a log_2_ fold change > 1 and an adjusted p-value < 0.05. Pseudobulk gene counts were aggregated across all cells, and differential expression analysis was performed using a negative binomial generalized linear model. P-values were determined by the Wald test and adjusted for multiple comparisons using the Benjamini-Hochberg method. The x-axis represents the log_2_ fold change in expression, while the y-axis shows the –log_10_ of the adjusted p-value.

(B) Ingenuity Pathway Analysis (IPA) identifying pathways upregulated in IL-9-treated compared with Mock-treated CAR T cells, shown as bar graphs. For this analysis, differentially expressed genes from the pseudobulk analysis with an adjusted p-value < 0.05 and fold change > 1.5 were included. Top pathways were identified using a z-score > 2, and pathways with a negative z-score were excluded from analysis.

(C) Illustration of CAR T cells transduced to express a PSMA-targeted CAR (Pbbz), dominant-negative TGF-βRII (dnTGF-βRII), and IL-9R.

(D) Transduction efficiency of human untransduced (UTD), dnTGF-βRII-Pbbz, or dnTGF-βRII-Pbbz-IL9R T cells analyzed by flow cytometry.

(E) *In vitro* production of human IL-9 (hIL-9) by PC3-PSMA cells treated with Ad-hIL9. Error bars indicate mean ± SD (n = 3).

(F) Experimental design schematic for xenograft prostate cancer model. SC, subcutaneous; IT, intratumoral; IV, intravenous; α-hIL10, neutralizing antibody targeting human IL-10.

(G)Concentration of human IL-10 in the serum analyzed via cytometric bead array on day D27 post-treatment (n = 10 for dnTGF-bRII-PBBz-IL9R 0.03E6 + Ad-hIL9, dnTGF-bRII-PBBz-IL9R 0.03E6, dnTGF-bRII-PBBz 0.03E6, and dnTGF-bRII-PBBz 1E6 cohorts; n = 8 for dnTGF-bRII-PBBz-IL9R 0.03E6 + Ad-hIL9 + α-hIL10 cohort).

(H) Tumor volume progression over time.

(I) Kaplan-Meier survival curves (data combined from donors ND623 and ND627).

(J) Count of infused T cells (hCD45+) in the blood, analyzed via flow cytometry on day 19 post-treatment (n = 10 for dnTGF-βRII-PBBz-IL9R 0.03E6 + Ad-hIL9, dnTGF-bRII-PBBz-IL9R 0.03E6, dnTGF-βRII-PBBz 0.03E6, and dnTGF-βRII-PBBz 1E6 cohorts; n = 8 for dnTGF-βRII-PBBz-IL9R 0.03E6 + Ad-hIL9 + α-hIL10 cohort).

Illustrations created with BioRender.com. Error bars indicate mean ± SD. Kruskal-Wallis one-way ANOVA was used for statistical analysis. For Kaplan-Meier survival curves, statistical significance was calculated using the log-rank Mantel-Cox test. ns (not significant) *P* > 0.05; \**P* ≤ 0.05; \*\**P* ≤ 0.01; \*\*\**P* ≤ 0.001; \*\*\*\**P* ≤ 0.0001.

**Supplemental Figure 6: Overview of UMAP clustering and CAR frequencies across treatments and donors, related to Figure 5**.

(A) Uniform manifold approximation and projection (UMAP) visualization of 70,097 cells from all donors and treatments, following data preprocessing. Cells are color-coded according to semi-supervised clustering annotations.

(B) Frequencies of CAR+ cells across all clusters, displayed after random down-sampling to match the donor with the lowest CAR+ frequency. Statistical comparisons between Mock and IL-9 treatments are provided, with Day 0 included as a baseline reference.

Cluster composition analysis by Welch’s t-test. ns (not significant) *P* > 0.05; \**P* ≤ 0.05; \*\**P* ≤ 0.01; \*\*\**P* ≤ 0.001; \*\*\*\**P* ≤ 0.0001.

**Supplemental Figure 7: Additional trajectories and gene expression heatmaps showing pseudotime-dependent changes along cellular trajectories, and STAT4 activity, related to Figures 6 and 7**.

(A) Two trajectories were identified but deemed non-informative.

Trajectory 5: Originating in the CD4+ naïve population and transitioning toward a CD4+ central memory phenotype. This trajectory was considered non-informative due to the absence of statistically significant genes between Mock and IL-9 treatments, as identified by the negative binomial generalized additive model (nb-GAM).

Trajectory 6: Initiating from a mixed naïve population of CD4+ and CD8+ T cells, traversing through CD4+ and CD8+ clusters, and culminating in a distinct subset of gamma-delta T cells. This trajectory was labeled non-informative as the pseudotime-mapped progression did not correspond with known biological phenotype transition states.

(C) Heatmaps were generated using genes identified by the nb-GAM model for the corresponding trajectory. Pseudotime values were divided into 40 bins (x-axis), with gene expression levels (y-axis) plotted across these pseudotime bins, highlighting temporal changes in gene activity.

(D) UMAP plots displaying STAT4 activity inferred using a univariate linear model (ULM). Transcription factor activity was determined using the Collection of Transcriptional Regulatory Interactions (CollecTRI) database. Interactions were weighted by activation or inhibition to assess transcription factor activity.

(E) STAT4 activity quantified for CAR+ cells in cluster 9 (CD8+ Exhausted). Statistical significance was assessed using a Kruskal-Wallis test.

## EXPERIMENTAL MODEL AND STUDY PARTICIPANT DETAILS

### Cell lines

The murine pancreatic ductal adenocarcinoma (PDA) cell line PDA7940b was derived from the Kras^LSL.G12D/+^p53^R172H/^+ (KPC) mouse pancreatic tumor model and was generously provided by Dr. Gregory Beatty from the University of Pennsylvania. PDA7940b cells were cultured in R10 media, comprising RPMI 1640 (Gibco) supplemented with 10% heat-inactivated fetal bovine serum (FBS, Seradigm), 2% 1 M HEPES buffer solution (Gibco), 1% 100X glutaMAX (Gibco), and 1% 10,000 U/mL penicillin and 10,000 μg/mL streptomycin (Gibco). Cells were routinely tested for mycoplasma contamination by the Department of Genetics at the University of Pennsylvania using the MycoAlert Mycoplasma Detection Kit (Lonza).

The human PDA cell line AsPC-1 and the human prostate cancer cell line PC3 were sourced from the American Type Culture Collection (ATCC). AsPC-1 mesothelin (MSLN) knock-out (KO) cells were generated by CRISPR-Cas9 technology, using Aldevron’s SpyFi™ Cas9 nuclease and sgRNA procured from Integrated DNA Technologies. The KO cells underwent three rounds of sorting using FACSAria™ Fusion Sorter (BD Biosciences). The PC3 cell line was genetically modified to express the PSMA protein, click beetle green luciferase, and green fluorescent protein (GFP) followed by purification, as previously described [58]. AsPC-1 WT and MSLN KO cells were cultured in in D20 medium, consisting of Dulbecco’s modified Eagle’s medium (DMEM) (1X, Gibco) supplemented with 20% heat-inactivated fetal bovine serum (FBS, Seradigm), 2% 1 M HEPES buffer solution (Gibco), 1% 100X glutaMAX (Gibco), and 1% 10,000 U/ml penicillin and 10,000 μg/ml streptomycin (Gibco). Modified PC3 cells were cultured in D10 medium, consisting of DMEM (1X, Gibco) supplemented with 10% FBS (Seradigm), 2% 1 M HEPES buffer solution (Gibco), and 1% 10,000 U/ml penicillin and 10,000 μg/ml streptomycin (Gibco). All cell lines were regularly authenticated by the University of Arizona Genetics Core and were routinely tested for mycoplasma contamination by the Department of Genetics at the University of Pennsylvania using the MycoAlert Mycoplasma Detection Kit (Lonza).

### Mice

All animal experiments were approved by the Institutional Animal Care and Use Committee (IACUC) at the University of Pennsylvania (protocol number: 804226), and procedures were carried out in compliance with federal and institutional IACUC guidelines. NOD/scid/IL2rγ^−/−^ (NSG) mice were initially obtained from Jackson Laboratories and subsequently bred at the Stem Cell & Xenograft Core (SCXC) at the University of Pennsylvania. For syngeneic mouse studies, C57BL/6 and B6.SJL-Ptprc^a^ Pepc^b^/BoyJ mice were sourced from Jackson Laboratories. Male (NSG) and female (NSG, C57BL/6 and B6.SJL-Ptprca Pepcb/BoyJ) mice, aged six to eight weeks, were used for *in vivo* experiments and maintained in pathogen-free conditions. Schematics of the mouse models employed are provided in both main text and supplementary figures.

Tumor sizes were measured one day prior to treatment to calculate initial tumor volumes, and mice were grouped into cohorts with comparable average tumor sizes. Animals with abnormally fast or slow tumor growth were excluded before treatment. Data were analyzed based on measurable outcomes without blinding. Mice underwent regular veterinary monitoring for signs of illness and were euthanized either at the end of the study or upon reaching pre-established IACUC rodent health endpoints. At no point did tumors exceed the maximum permissible diameter of 2 cm, as set by the institutional review board.

### Human Samples

Healthy donor primary T lymphocytes were provided by the University of Pennsylvania Human Immunology Core. Samples are de-identified for compliance with HIPAA rules. Donor sex and age is shown below: ND552 (female, age 29), ND541 (female, age 31), TMP489 (female, age 23), ND627 (male, age 30), ND623 (female, age 25).

Frozen serum samples were collected from pancreatic cancer patients who enrolled in a clinical trial of human mesothelin-targeted (M5) CAR T cell therapy. Patients enrolled in this trial were adults aged 18 years or older with histologically confirmed unresectable or metastatic pancreatic adenocarcinoma. All patients gave informed consent in accordance with the Declaration of Helsinki. This study was registered at ClinicalTrials.gov (identifier NCT03323944).

## METHOD DETAILS

### General cell culture

The murine pancreatic ductal adenocarcinoma (PDA) cell line PDA7940b was derived from the Kras^LSL.G12D/+^p53^R172H/+^ (KPC) mouse pancreatic tumor model and was generously provided by Dr. Gregory Beatty from the University of Pennsylvania.

PDA7940b cells were cultured in R10 media, comprising RPMI 1640 (Gibco) supplemented with 10% heat-inactivated fetal bovine serum (FBS, Seradigm), 2% 1 M HEPES buffer solution (Gibco), 1% 100X GlutaMAX (Gibco), and 1% 10,000 U/ml penicillin and 10,000 μg/ml streptomycin (Gibco). Cells were routinely tested for mycoplasma contamination by the Department of Genetics at the University of Pennsylvania using the MycoAlert Mycoplasma Detection Kit (Lonza).

The human PDA cell line AsPC-1 and the human prostate cancer cell line PC3 were sourced from the American Type Culture Collection (ATCC). AsPC-1 MSLN knock-out (KO) cells were generated by CRISPR-Cas9 technology, using Aldevron’s SpyFi™ Cas9 nuclease and sgRNA procured from Integrated DNA Technologies. The KO cells underwent three rounds of sorting using FACSAria™ Fusion Sorter (BD Biosciences). The PC3 cell line was genetically modified to express the PSMA protein, click beetle green luciferase, and green fluorescent protein (GFP) followed by purification, as previously described [58]. AsPC-1 WT and MSLN KO cells were cultured in D20 medium, consisting of Dulbecco’s modified Eagle’s medium (DMEM) (1X, Gibco) supplemented with 20% heat-inactivated fetal bovine serum (FBS, Seradigm), 2% 1 M HEPES buffer solution (Gibco), 1% 100X GlutaMAX (Gibco), and 1% 10,000 U/ml penicillin and 10,000 μg/ml streptomycin (Gibco). Modified PC3 cells were cultured in D10 medium, consisting of DMEM (1X, Gibco) supplemented with 10% FBS (Seradigm), 2% 1 M HEPES buffer solution (Gibco), and 1% 10,000 U/ml penicillin and 10,000 μg/ml streptomycin (Gibco). All cell lines were regularly authenticated by the University of Arizona Genetics Core and were routinely tested for mycoplasma contamination by the Department of Genetics at the University of Pennsylvania using the MycoAlert Mycoplasma Detection Kit (Lonza).

### Murine CAR T cell production

Retroviral transduction of mouse CAR T cells was previously described [59]. Briefly, spleens from CD45.1+ B6.SJL-^Ptprca Pepcb^/BoyJ mice were harvested, and T cells were purified with mouse T cell isolation beads (STEMCELL Technologies). T cells were activated with mouse CD3/CD28 Dynabeads (Gibco) at a bead-to-T cell ratio of 2:1. After 48 hours, T cells were transduced with retroviral vectors encoding the anti-mesothelin CAR (A03 CAR) or A03 CAR and IL-9R on recombinant human fibronectin-coated plates (RetroNectin, Takara Bio). Cells were cultured in mouse T cell media consisting of RPMI 1640 (Gibco) supplemented with 10% heat-inactivated FBS (Seradigm), 1% 100X GlutaMAX (Gibco), 1% 10,000 U/mL penicillin + 10,000 ug/mL streptomycin (Gibco), 1X 100 mM sodium pyruvate (Gibco), and 50 uM β-mercaptoethanol (Gibco) supplemented with recombinant mouse IL-2 (Abcam) (50 U/mL). At day five after stimulation, mouse CAR T cells were harvested, de-beaded, and used for *in vitro*/*in vivo* experiments.

### Human CAR T cell production

Lentiviral vectors and human CAR T cells were generated as previously described [60]. Human CAR T cells were produced using T cells from healthy donors obtained from the Human Immunology Core (HIC) at the University of Pennsylvania. For the production of CAR T cells targeting mesothelin, freshly isolated CD4+ and CD8+ T cells were mixed in a 1:1 ratio and activated with human CD3/CD28 Dynabeads (Gibco) at a bead-to-cell ratio of 3:1. Twenty-four hours later, T cells were transduced with lentiviral vectors encoding the anti-mesothelin M5 CAR, M5 CAR and IL-9R, dominant negative TGF-βRII (dnTGF-βRII) and anti-PSMA CAR (Pbbz), or dn-TGFβRII-PBBz and IL-9R, using a multiplicity of infection (MOI) of 3. On day five, the CD3/CD28 beads were removed from the culture. T cells were maintained in R10 media and cultured at a density of 0.8 x 10^6 cells per mL until they reached a resting state, as determined by cell volume of ∼350 fL using a Multisizer 4 Coulter Counter (Beckman). For cryopreservation of T cells, freezing media consisting of 90% heat-inactivated FBS (Seradigm) and 10% dimethyl sulfoxide (DMSO, Sigma-Aldrich) was used.

### Phosphoflow signaling assay

Mouse T cells were starved overnight in mouse T cell media without IL-2. Human T cells were starved for 4 hours in RPMI 1640 (Gibco) containing 0.1% FBS (Seradigm). Cells were then stimulated by adding recombinant mouse IL-9 or animal-free human IL-9 (Peprotech) and incubated for 20 minutes at 37°C. To terminate the reaction, cells were fixed with 1.5% formaldehyde (Cell Signaling Technology) for 15 minutes at room temperature with agitation. After fixation, cells were washed with 1X DPBS (Gibco) and then permeabilized using ice-cold 100% methanol (Cell Signaling Technology) for 1 hour on ice. Post-permeabilization, cells were washed with buffer consisting of 1X DPBS (Gibco), 0.5% heat-inactivated FBS (Seradigm), 2.5 mM EDTA (Invitrogen) and 1% 10,000 U/mL penicillin + 10,000 µg/mL streptomycin (Gibco). Cells were subsequently stained for 1 hour at 4°C in the dark with antibodies targeting phospho-STAT1 (Tyr701) (Cell signaling, 8062S), phospho-STAT3 (pY705) (BD Biosciences, 557815) and phospho-STAT5 (pY694) (BD Biosciences, 560117). The stained cells were washed again and analyzed using an LSRFortessa cytometer (BD Biosciences). The data were represented as log fold change of mean fluorescence intensity (MFI), and the results were fitted to a log(agonist) versus dose-response model using Prism 10 software (GraphPad).

### CAR T cell *in vitro* dysfunction model

AsPC-1 cells were routinely seeded in 12-well plates at a density of 0.1 x 10^6 cells per well the day before introducing T cells. CAR T cells were thawed in R10 medium and allowed to rest overnight. After 24 hours, T cells were counted, and transduction efficiency was assessed by flow cytometry. Cell viability was determined using a Countess automated cell counter (ThermoFisher Scientific). 0.1 x 10^6 viable CAR T cells per well were then added to the AsPC-1-coated plates. After a 2-day co-culture period, the cells were thoroughly resuspended by frequent pipetting. The resulting cell suspension was centrifuged, and the supernatant (conditioned media) was collected and filtered using a 0.45 μm filter (Corning). The cell pellet was resuspended in a mixture of conditioned media and fresh R10 medium in equal proportions. The T cell suspension was then transferred back to the AsPC-1-coated plates for continued co-culture. This process was repeated every 2 days for a total duration of 10 days. For the IL-9-treated group, animal-free human IL-9 (10 nM, Peprotech) was added starting at the beginning of the assay and every other day. Flow cytometry analysis was performed on Day 0 and 10 for T cell immunophenotyping. For scRNA-seq analysis, live, EpCAM-, CD45+ T cells were sorted as described below.

### Cytokine production assays

For cytokine secretion analysis with mouse T cells, cells were incubated in mouse T cell media with or without murine IL-9 (100 nM, Peprotech) for 48 hours, and the resulting cell supernatants were submitted to the Human Immunology Core (HIC) at the University of Pennsylvania for analysis using a Luminex assay. The Th1/Th2/Th9/Th17/Th22/Treg Cytokine 17-Plex Mouse ProcartaPlex™ Panel (Invitrogen) was used to detect mouse cytokines.

For human T cell cytokine secretion analysis, CAR T cells were thawed and incubated in R10 media supplemented with animal-free human IL-9 (10 nM, Peprotech) or without IL-9 (Mock) for 24 hours. The supernatants were then collected and analyzed using the Human Th1/Th2/Th17 Cytokine CBA Kit (BD Biosciences), following the manufacturer’s instructions. Sample data were acquired using a FACSymphony A5 flow cytometer (BD Biosciences) equipped with FACSDiva software, and the results were processed using BD CBA Analysis Software (BD Biosciences).

For cytokine secretion analysis following the *in vitro* dysfunction model, human CAR T cells were co-cultured overnight with AsPC-1 cells at a 1:1 effector-to-target cell ratio, or with media as a control. Cytokine concentrations in the supernatants were determined using the Human Th1/Th2/Th17 Cytokine CBA Kit (BD Biosciences), as per the manufacturer’s instructions. Sample data were acquired using a FACSymphony A5 flow cytometer (BD Biosciences) equipped with FACSDiva software, and the results were processed using BD CBA Analysis Software (BD Biosciences).

To measure IL-9 production following Ad-mIL9 or Ad-hIL9 infection of tumor cells *in vitro*, PDA7940b or AsPC-1 cells were cultured with Ad-mIL9 or Ad-hIL9 (100 VP/cell) respectively, or with media as a control (Mock). Supernatants were collected at various timepoints, and IL-9 concentrations were assessed using murine or human IL-9 ELISA kits (Abcam). Absorbance readings were obtained using a Synergy H4 plate reader (BioTek).

For the detection of IL-9 in human serum, samples from healthy donors, provided by the Human Immunology Core (HIC) at the University of Pennsylvania, or from patients in clinical trial UPCC14217 (NCT03323944, see Table S1 for sample IDs) provided by the Translational and Correlative Studies Laboratory (TCSL) were analyzed. IL-9 concentrations were determined using a human IL-9 ELISA kit (Abcam, minimal detectable dose 2.3 pg/mL), and the readings were taken on a Synergy H4 plate reader (BioTek).

### *In vitro* cytotoxicity assays

*In vitro* analysis of CAR T cell killing was performed using a real-time, impedance-based assay with xCELLigence Real-Time Cell Analyzer System (Agilent). Briefly, 10,000 PDA7940b cells, AsPC-1 WT cells or AsPC-1 MSLN KO cells were seeded to a 96-well E-plate. After 24 hours, CAR T cells were added to the wells in different effector-to-target cell (E:T) ratios as indicated for each assay. For experiments with murine CAR T cells, cells were incubated in mouse T cell media with or without IL-9 (100 nM) for 48 hours prior to addition into xCELLigence plates. Tumor killing was monitored every 20 minutes over a total of 5 days. For data analysis, cell counts were normalized to the time point of CAR T cell addition.

### Xenograft animal models

For xenograft flank tumor models, AsPC-1 or PC3-PSMA cells were utilized to establish tumors. A total of 2 x 10^6 AsPC-1 cells or 1 x 10^6 PC3-PSMA cells, suspended in 100 μL of a 1:1 mix of Matrigel (Corning) and 1X DPBS (Gibco), were implanted subcutaneously into the right flank of NSG mice. Tumor growth was monitored until it reached a volume of approximately 150-200 mm³. After 15 days, mice received an intratumoral (IT) injection of 1 x 10^9 viral particles (VP) of Ad-hIL9 (Vector Biolabs) in 50 μL 1X DPBS. The following day, mice were treated with either 0.03 x 10^6 or 1 x 10^6 CAR T cells, depending on their assigned treatment group. A second dose of Ad-hIL9 (1 x 10^9 VP in 50 μL) was administered via IT injection three days later. Control groups received either Ad-hIL9 alone, CAR T cells alone, or no treatment. For groups not receiving Ad-hIL9, an IT injection of 1X DPBS was given as a control. For the tumor re-challenge experiment, mice were implanted with 2 x 10^6 AsPC-1 cells by SC injection into the left flank, and treatment-naïve mice were used as a control. In the xenograft prostate cancer model, one cage of mice (from the group treated with CAR T 0.03E6) was flooded on day 67, leading to the removal of this cage from the survival data analysis.

In antibody neutralization experiments, mice received intraperitoneal (IP) injections of either 300 μg of anti-murine IL-9 antibody (BioXcell, BE0181), 300 μg of isotype control antibody (BioXcell, BE0085), or 500 μg of anti-human IL-10 antibody (BioXcell, BE0441) diluted in 1X DPBS starting on the day of the first IT injection and continuing every 3 days.

Tumor size and mouse body weight were measured at least once a week, with values normalized to the day of the first IT injection for analysis presented in the main and supplementary figures. In a separate experiment, NSG mice were sacrificed on day 25 post-treatment for staining of human CD45+ cells in tumor tissues and peripheral blood. Health assessments were conducted following IACUC’s body condition scoring (BCS) guidelines.

### Syngeneic animal model

C57BL/6 mice were subcutaneously injected with 5 x 10^5 PDA7940b tumor cells in 100 μL of 1X DPBS (Gibco) into the right flank. On day six, mice underwent lymphodepletion via intraperitoneal injection of cyclophosphamide (120 mg/kg, Sigma-Aldrich). The following day, when tumors reached a size of approximately 50-100 mm³, an intratumoral (IT) injection of Ad-mIL9 (Vector Biolabs) at a dose of 1 x 10^9 VP in 50 μL 1X DPBS was administered. On the next day, 5 x 10^6 viable CAR T cells were injected intravenously (IV) in 150 μL of PBS. Three days later, a second IT dose of Ad-mIL9 at the same dose was delivered. Control groups received either CAR T cells or Ad-mIL9 alone. For groups not receiving Ad-mIL9, an IT injection of PBS was administered as a control.

Tumor size and body weight were measured at least twice weekly using calipers, and data were normalized to the day of the first IT injection in the main and supplementary figures where applicable. In specific experiments, mice were sacrificed on day 5 or 6 post-treatment for several analyses: (i) flow cytometry to assess the numbers of CAR T cells and other immune cells in tumors, spleens, and peripheral blood, (ii) pathological assessment, immunohistochemistry (IHC), and RNA in situ hybridization (RNA ISH) of tumor tissues, and (iii) cytokine analysis from mouse serum. Health monitoring was conducted according to the Institutional Animal Care and Use Committee (IACUC) body condition scoring (BCS) guidelines.

### Caliper measurements of subcutaneous tumors

Tumor size was measured using calipers, and volumes were calculated with the formula: volume = (length in mm × width² in mm) / 2. Tumor size values were normalized to the day of the first IT injection in the main and supplementary figures where indicated.

### Preparation of single-cell suspensions from tumors and spleens

Tumors and spleens were collected and processed into single-cell suspensions as previously detailed [20]. Briefly, after harvesting, tumor tissue was finely chopped into small pieces (3-5 mm) using a scalpel. The fragments were digested in DMEM (Gibco) containing Collagenase/Hyaluronidase (STEMCELL Technologies) and DNase I (1 mg/mL, STEMCELL Technologies) at 37°C while shaking at 200 rpm for 30 minutes. To remove red blood cells, ACK Lysis Buffer (Life Technologies) was applied. Cells were then stained for flow cytometry analysis. For spleens, tissue was gently mashed using the flat end of a syringe plunger, and red blood cells were lysed with ACK Lysis Buffer for subsequent flow cytometry staining and analysis.

### Peripheral blood (CAR) T cell analysis

Peripheral blood was collected from C57BL/6 or NSG mice via cardiac puncture for terminal studies or retroorbital bleeding in overall survival studies. Samples were stained and quantified using TruCount tubes (BD Biosciences) as per the manufacturer’s guidelines. Data acquisition was performed on an LSRFortessa cytometer (BD Biosciences) equipped with FACSDiva software (BD Biosciences). Data were processed and analyzed using FlowJo version 10 software (BD Biosciences).

### Cytokine analysis from mouse serum

To collect serum from C57BL/6 mice, whole blood was obtained via cardiac puncture, allowed to clot at room temperature for 20 minutes, and then centrifuged to separate the serum. Mouse serum was then sent to the Human Immunology Core (HIC) at the University of Pennsylvania for cytokine analysis using a Luminex assay. The Th1/Th2/Th9/Th17/Th22/Treg Cytokine 17-Plex Mouse ProcartaPlex™ Panel (Invitrogen) was employed to detect various mouse cytokines.

For cytokine analysis in NSG mice, whole blood was obtained via retroorbital bleeding and serum was collected. Serum samples were analyzed using the human IL-9 CBA kit, murine IL-9 CBA kit, or the human Th1/Th2/Th17 Cytokine CBA kit (BD Biosciences), following the manufacturer’s instructions. Data acquisition was performed using the FACSymphony A5 flow cytometer (BD Biosciences) with FACSDiva software, and results were analyzed with the BD CBA Analysis Software (BD Biosciences).

### Immunohistochemistry (IHC) staining, scanning and image analysis

Tumors of C57BL/6 mice were harvested and prepared for standard IHC analysis by the Comparative Pathology Core (CPC) at the University of Pennsylvania School of Veterinary Medicine. Formalin fixed tissues were routinely processed for paraffin embedding, sectioning, and staining for hematoxylin and eosin (H&E). For immunohistochemistry, 5 μm thick sections were mounted on ProbeOn^TM^ slides (Thermo Fisher Scientific) and stained using a Leica BOND RXm automated platform combined with the Bond Polymer Refine Detection kit (Leica #DS9800). Briefly, after dewaxing and rehydration, sections were pretreated with the epitope retrieval BOND ER2 high pH buffer (Leica #AR9640) for 20 minutes at 98°C. Endogenous peroxidase was inactivated with 3% hydrogen peroxide for 10 minutes at room temperature (RT). Nonspecific tissue-antibody interactions were blocked with Leica PowerVision IHC/ISH Super Blocking solution (PV6122; Leica Biosystems, Deer Park, IL) for 30 minutes at RT. The same blocking solution also served as diluent for the primary antibodies. A battery of primary antibodies for immune cells were used and included CD3 (Bio-rad, MCA1477T, rat monoclonal antibody, 1/600), FoxP3 (CST, 12653, rabbit monoclonal antibody, 1/300), K1rb1c/CD161c (CST, 39197, rabbit monoclonal antibody, 1/500NK), Pax5 (CST, 12709, rabbit monoclonal antibody, 1/100), F4/80 (CST, 70076, rabbit monoclonal antibody, 1/1000), CD11c (CST, 97585, rabbit monoclonal antibody, 1/150), Ly6G (CST, 87048, rabbit monoclonal antibody, 1/100), tryptase (CST, 19523, rabbit monoclonal antibody, 1/200), and PRG2 (Novus Bio, NBP3-04784, rabbit polyclonal antibody, 1/300). A biotin-free polymeric IHC detection system consisting of horseradish peroxidase (HRP)-conjugated antirabbit or antirat IgG was then applied for 25 minutes at RT. Immunoreactivity was revealed with the diaminobenzidine (DAB) chromogen reaction. Slides were counterstained in hematoxylin, dehydrated in an ethanol series, cleared in xylene, and permanently mounted with a resinous mounting medium (Thermo Scientific ClearVue coverslipper). Slides were then scanned at 20x magnification using the Leica Versa 200 whole slide scanner (Leica Biosystem), visualized and analyzed using the Aperio ImageScope software (Leica Biosystem). For the analysis, nuclear and cytoplasmic cell counting algorithms were used for FoxP3 and PAX5, and for K1RB1C and tryptase, respectively. For cell counting algorithms (PAX5, K1RB1C, FoxP3 and tryptase), the number of positive cells was normalized to the area of analysis on each slide. Positive pixel count algorithms were used for the quantification of area positivity for CD11c, F4/80, Ly6G, and PRG2. For these markers, the positivity was calculated by dividing the number of positive pixels by the total number of pixels in the area of analysis.

### RNA in situ hybridization (RNA ISH)

RNA ISH to visualize RNA molecules in individual cells of formalin-fixed, paraffin-embedded tumor tissues was performed according to manufacturer’s instructions (RNAscope, ACDBio). The following probes were used: Mm-Msln (CAT: 443241, ACDBio), Mm-Cd3e-C2 (CAT: 314721-C2, ACDBio), vMSGV-C3 (CAT:1140971-C3, ACDBio), 4-plex Negative Control Probe (CAT: 321831, ACDBio). The 4-plex Negative Control Probe was used to confirm that the samples were free of non-specific signal or interfering substances that would confound interpretation and quantification. The slides were scanned at 20x or 40x magnification using the Leica Versa 200 whole slide scanner (Leica Biosystem), visualized and analyzed using the Aperio ImageScope software (Leica Biosystem).

### Flow cytometry and sorting

For flow cytometry, cells were stained in fluorescence-activated cell sorting (FACS) buffer consisting of 1X DPBS (Gibco), 0.5% heat-inactivated FBS (Seradigm), 2.5 mM EDTA (Invitrogen) and 1% 10,000 U/mL penicillin + 10,000 ug/mL streptomycin (Gibco). For sorting, cells were stained in FACS buffer with 100mg/mL DNase (Sigma-Aldrich). Antibodies specific for human CD4 (300558), CD8 (300934), CD45RA (304114), IL-9R (310404), EpCAM (324206), PD-1 (329920), TIM3 (345016), CD27 (356412) and CD45 (368524/304032) and for mouse CD3 (100229), CD11b (101242), CD44 (103030), CD45 (103128), CD62L (104406), CD86 (105020), MHCII (107636), NK1.1 (108748), CD45.1 (110741), CD19 (115543), CD11c (117318), F4/80 (123128), Ly6G (127626), Ly6C (128032), CD206 (141706), CD95 (152604) and IL-9R (158704) were purchased from Biolegend. Antibodies specific for phospho-STAT3 (pY705) (557815) and phospho-STAT5 (pY694) (560117) were purchased from BD Biosciences. An antibody specific for Phospho-STAT1 (Tyr701) (8062S) was purchased from Cell Signaling. An antibody specific for human LAG3 (61-2239-42) was purchased from eBioscience. An antibody specific for anti-human TGF-beta RII (FAB2411A) was purchased from R&D systems. M5 CAR and A03 CAR expression was assessed using biotinylated goat anti-human IgG F(ab’)_2_ (Jackson ImmunoResearch, 109–066-006) followed by streptavidin (PE-or APC-conjugated, Biolegend). Pbbz expression was assessed using biotinylated goat Anti-Mouse IgG F(ab’)₂ (Jackson ImmunoResearch, 115-066-072). Live/dead cell staining was performed using a Live/Dead Aqua (Life Technologies) or Zombie NIR (Biolegend) Fixable Viability Kits following manufacturer’s protocol. TruStain FcX (anti-mouse CD16/32) Antibody (CAT: 101320; BioLegend) or Human TruStain FcX Fc Receptor Blocking Solution (CAT: 422302; BioLegend) were used prior to staining of single cell suspensions of mouse or human tumors and mouse spleens. CountBright™ Absolute Counting Beads (ThermoFisher) were used as an internal standard according to the manufacturer’s instructions to calculate absolute cell counts in cell suspensions. Samples were acquired on an LSRII Fortessa Cytometer (BD Biosciences) or FACSymphony™ A5 SE Cell Analyzer (BD Biosciences) and analyzed with FlowJo v10 software (FlowJo, LLC). Sorting was performed using a FACSAria™ Fusion Sorter (BD Biosciences).

### Single-cell RNA sequencing (scRNA-seq)

scRNA-seq was performed by the Genomics Facility at the Wistar Institute. Single cell droplets were generated using the Chromium Next GEM single cell 5’ kit v2 Dual Index kit (10x Genomics, Pleasanton, CA). cDNA synthesis and amplification, library preparation and indexing were performed using the 10x Genomics Library Preparation kit (10x Genomics), according to manufacturer’s instructions. Overall library size was determined using the Agilent TapeStation 4200 with the High Sensitivity DNA 5000 ScreenTape (Agilent, Santa Clara, CA) and libraries were quantitated using KAPA real-time PCR (Roche, Wilmington, MA). A total of 9 samples generating 9 libraries were pooled and sequenced on the NextSeq 2000 (Illumina, San Diego, CA) using 2 P3 300 cycle kits (Illumina) resulting in 250 million reads per sample. Run configuration was paired-end with the following parameters: 26 base pair (read1) x 8 base pair (index) x 281 base pair (read2).

### scRNA-seq data quality control and preprocessing

Sequencing quality control was conducted with FASTQC (v 0.11.9) and sources of contamination were detected with FASTQ Screen (v 0.14.0). Alignment and quantitation were conducted with CellRanger v7.2.0 using a custom reference genome based on GRCh38 2022 with our CAR sequence inserted. GEM Selection was completed by running the scGEM2Cellr package. Quality control and pre-processing was conducted with the scater package [61]. Doublets were removed using the scDblFinder package [62]. High mitochondrial cells were removed using a linear model of mitochondrial transcript percent by the total genes detected per cell [63]. Genes that were not found in at least 5 cells were discarded. Non-donor-specific variation in the expression of T cell receptor beta variable (TRBV) transcripts induced separation during dimensionality reduction, which was unrelated to the cell types of interest in our study. To address this, we excluded TRAV and TRBV genes from the principal component analysis (PCA) used to generate the Uniform Manifold Approximation and Projection (UMAP). However, these genes were retained for all downstream analyses, including differential gene expression analysis and gene set enrichment analysis.

### scRNA-seq data analysis

Data analysis was conducted using the standard Bioconductor analysis approach [64]. Batch effects were assessed using the scater package and addressed using Harmony integration [65, 66]. Data normalization was performed using the variance-stabilizing transformation model in sctransform v2 [67]. To focus on biological questions, non-biological phenotype clusters caused by T cell variable region transcripts were removed prior to dimensional reduction, which was performed using Uniform Manifold Approximation and Projection (UMAP) with 10 nearest neighbors and a minimum spanning distance of 0.5 [68].

Cluster stability was investigated using adjusted Rand Index and normalized mutual information with the Dune package [69]. Clusters were annotated by scoring cells against the Human Cell Atlas, Pan-Cancer T cell Atlas, and prior publications [24, 70, 71]. Cell scoring was achieved by calculating the area under the curve for each signature, and clusters were labeled based on a guilt-by-association approach from the cell labels in each cluster [72]. Cluster composition was compared using Welch’s t-test.

Differential gene expression was conducted using the MAST package [73]. Gene Set Enrichment Analysis (GSEA) was performed on statistically significant genes using the following databases: MSigDB Hallmark 2020, KEGG 2021 Human, Reactome 2022, GO Biological Process 2023, GO Cellular Component 2023, and GO Molecular Function 2023, implemented with the R package EnrichR [74].

For pseudobulk analysis, gene counts were calculated by aggregating the expression data from all cells within each group. Differential expression analysis was conducted using a negative binomial generalized linear model, with p-values calculated via the Wald test and adjusted for multiple comparisons using the Benjamini-Hochberg method. Ingenuity Pathway Analysis (IPA, Qiagen Inc.) was utilized to identify differential canonical pathways. For this analysis, differentially expressed genes from the pseudobulk analysis with an adjusted p-value < 0.05 and fold change > 1.5 were included. Top pathways were identified using a z-score > 2, and pathways with a negative z-score were excluded from analysis.

Transcription factor (TF) analysis was performed using the TRRUST v2 database, following a process analogous to GSEA. To validate the most enriched and downregulated TF pathways, we conducted upstream regulator analysis with IPA. Differentially expressed genes from CD8+ or CD4+ CAR+ cells served as input, with a cutoff of ±1 log_2_ fold change. The top five TFs were selected based on concordance between TRRUST and IPA results, excluding those with z-scores below ±0.05. The activity of TFs was inferred using a univariate linear model (ULM). This approach utilized the Collection of Transcriptional Regulatory Interactions (CollecTRI) which compiles signed TF-target gene interactions from 12 different resources [75]. The ULM was applied to score the activity of each TF in individual cells.

Trajectory inference was conducted using the slingshot package [33]. Trajectory comparisons were analyzed using the tradeSeq workflow, and genes were identified by fitting a generalized additive model (GAM) for each gene [76]. Donor CAR+ frequency between conditions of trajectories were determined by Fisher’s repeated measures t-test. RNA velocity analysis was performed using the scVelo package. Unless otherwise specified, all scRNA-seq analyses were conducted using Python 3.13 or R 4.3.

### Statistical analysis

Statistical analysis was performed with Prism version 10 (GraphPad). Each figure legend denotes the statistical test used. Survival curves were drawn using the Kaplan-Meier method and the differences of 2 curves were compared with the log-rank Mantel-Cox test. Comparisons between two groups were performed using a two-tailed unpaired Student’s t-test. Kruskal-Wallis one-way analysis of variance (ANOVA) was used to compare 3 or more groups. Statistical significance between multiple groups of two variables was assessed by two-way ANOVA with post-hoc tests. For all figures, ns indicates *P* > 0.05 (non-significant), * indicates *P* ≤ 0.05, ** indicates *P* ≤ 0.01, *** indicates *P* ≤ 0.001, and **** indicates *P* ≤ 0.0001. Graphs were created by Prism version 10 (GraphPad), Adobe Illustrator (Adobe), and BioRender.com. All *in vitro* data presented are representative of at least two biological replicates. scRNA-seq was performed in three biological replicates (three donors). *In vivo* animal studies were performed using two biological replicates, independently.

## SUPPLEMENTAL TABLE TITLES

Table S1. ELISA quantification of IL-9 levels in serum from clinical trial patients and normal donors, related to Figures 2 and S2

Table S2. Full list of DEGs between IL-9-treated and Mock-treated CAR+ cells from pseudobulk analysis, related to Figures 4 and S5A

Table S3. Full list of IPA pathways enriched in DEGs between IL-9-treated and Mock-treated CAR+ cells, from pseudobulk analysis, related to Figures 4 and S5B

Table S4. Full list of DEGs between IL-9-treated and Mock-treated CD8+ and CD4+ CAR+ cells from the scRNA-seq experiment, related to Figure 5D

Table S5. Full list of GSEA pathways enriched in IL-9-treated compared with Mock-treated CD8+ and CD4+ CAR+ cells from the scRNA-seq experiment, related to Figure 5E

Table S6. Full list of transcription factor (TF) pathways enriched in IL-9-treated compared with Mock-treated CD8+ and CD4+ CAR+ cells analyzed using TRRUST database, related to Figure 7C

Table S7. Full list of upstream regulators predicted by IPA (upstream regulator analysis) using DEGs between IL-9-treated and Mock-treated CD8+ and CD4+ CAR+ cells, related to Figure 7C

