## Supplemental figures for "IL-9 signaling redirects CAR T cell fate toward CD8+ memory and CD4+ cycling states, enhancing anti-tumor efficacy"

Figure S1: IL-9 signaling enhances CAR T cell infiltration into tumor and spleen without markedly altering the TME, related to Figure 1.

Figure S2: Murine IL-9 activates IL-9R in human CAR-IL9R T cells without exogenous addition of IL-9, related to Figure 2.

Figure S3: IL-9-signaling induces a central memory phenotype and reduces co-inhibitory receptor expression in CAR T cells, related to Figure 3.

Figure S4: IL-9 signaling enhances CAR T cell effector function and promotes a central memory phenotype after repeated antigen stimulation across multiple donors, related to Figure 4.

Figure S5: IL-9-driven enhancement of CAR T cell antitumor efficacy in a prostate cancer xenograft model is independent of IL-10 signaling, related to Figure 4.

Figure S6: Overview of UMAP clustering and CAR frequencies across treatments and donors, related to Figure 5.

Figure S7: Additional trajectories and gene expression heatmaps showing pseudotime-dependent changes along cellular trajectories, and STAT4 activity, related to Figures 6 and 7.

### Supplemental Tables

Table S1. ELISA quantification of IL-9 levels in serum from clinical trial patients and normal donors, related to Figures 2 and S2

Table S2. Full list of DEGs between IL-9-treated and Mock-treated CAR+ cells from pseudobulk analysis, related to Figures 4 and S5A

Table S3. Full list of IPA pathways enriched in DEGs between IL-9-treated and Mock-treated CAR+ cells, from pseudobulk analysis, related to Figures 4 and S5B

Table S4. Full list of DEGs between IL-9-treated and Mock-treated CD8+ and CD4+ CAR+ cells from the scRNA-seq experiment, related to Figure 5D

Table S5. Full list of GSEA pathways enriched in IL-9-treated compared with Mock-treated CD8+ and CD4+ CAR+ cells from the scRNA-seq experiment, related to Figure 5E

Table S6. Full list of transcription factor (TF) pathways enriched in IL-9-treated compared with Mock-treated CD8+ and CD4+ CAR+ cells analyzed using TRRUST database, related to Figure 7C

Table S7. Full list of upstream regulators predicted by IPA (upstream regulator analysis) using DEGs between IL-9-treated and Mock-treated CD8+ and CD4+ CAR+ cells, related to Figure 7C

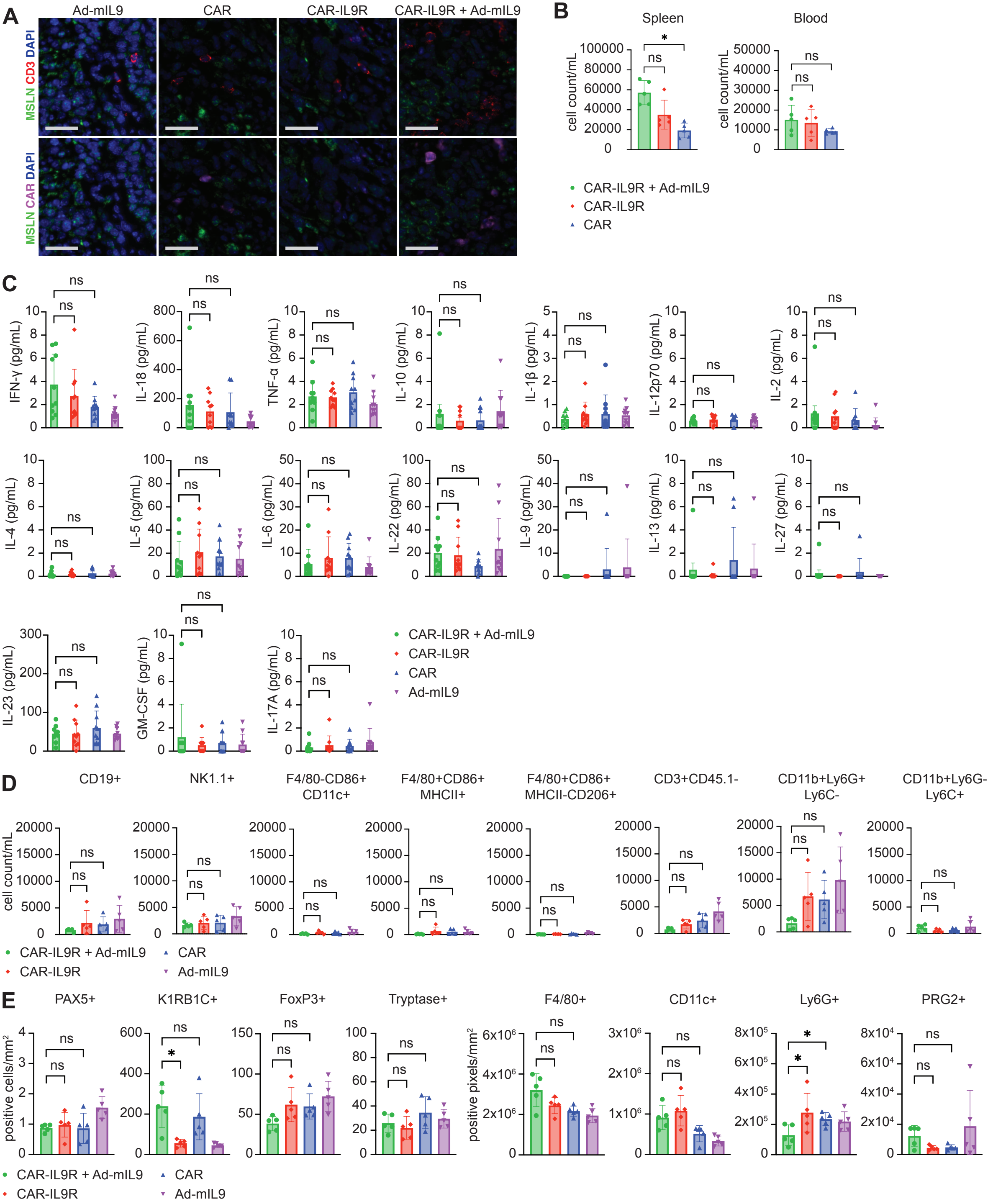

A
Ad-mIL9
CAR
CAR-IL9R
CAR-IL9R + Ad-mIL9
MSLN CD3 DAPI
MSLN CAR DAPI
B
Spleen
Blood
cell count/mL
CAR-IL9R + Ad-mIL9
CAR-IL9R
CAR
C
IFN-γ (pg/mL)
IL-18 (pg/mL)
TNF-α (pg/mL)
IL-10 (pg/mL)
IL-1β (pg/mL)
IL-12p70 (pg/mL)
IL-2 (pg/mL)
IL-4 (pg/mL)
IL-5 (pg/mL)
IL-6 (pg/mL)
IL-22 (pg/mL)
IL-9 (pg/mL)
IL-13 (pg/mL)
IL-27 (pg/mL)
IL-23 (pg/mL)
GM-CSF (pg/mL)
IL-17A (pg/mL)
CAR-IL9R + Ad-mIL9
CAR-IL9R
CAR
Ad-mIL9
D
CD19+
NK1.1+
F4/80-CD86+ CD11c+
F4/80+CD86+ MHCII+
F4/80+CD86+ MHCII-CD206+
CD3+CD45.1-
CD11b+Ly6G+ Ly6C-
CD11b+Ly6G- Ly6C+
cell count/mL
CAR-IL9R + Ad-mIL9
CAR-IL9R
CAR
Ad-mIL9
E
PAX5+
K1RB1C+
FoxP3+
Tryptase+
F4/80+
CD11c+
Ly6G+
PRG2+
positive cells/mm²
positive pixels/mm²
CAR-IL9R + Ad-mIL9
CAR-IL9R
CAR
Ad-mIL9

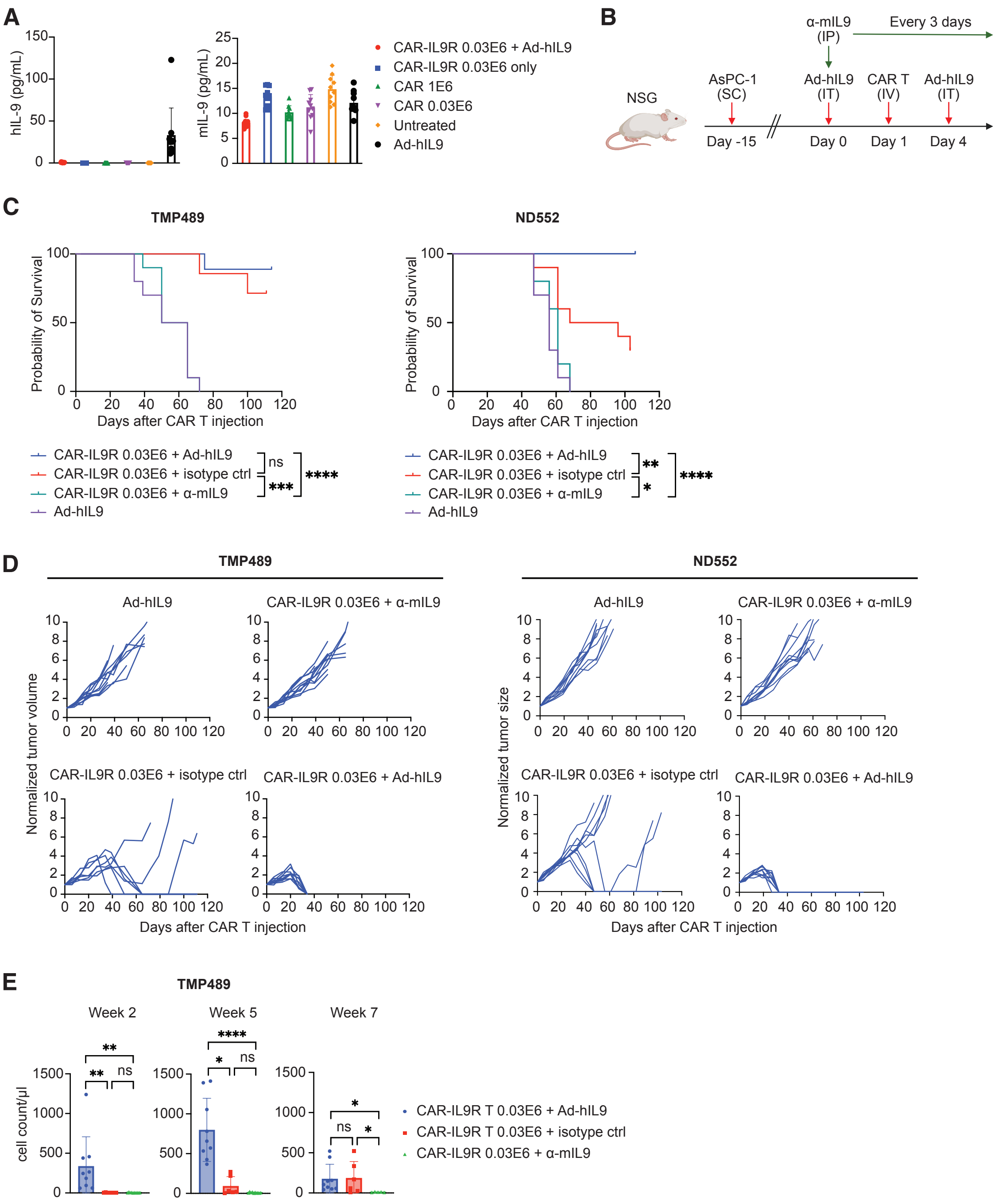
A
hIL-9 (pg/mL)
mIL-9 (pg/mL)
CAR-IL9R 0.03E6 + Ad-hIL9
CAR-IL9R 0.03E6 only
CAR 1E6
CAR 0.03E6
Untreated
Ad-hIL9
B
NSG
α-mIL9 (IP)
Every 3 days
AsPC-1 (SC)
Ad-hIL9 (IT)
CAR T (IV)
Ad-hIL9 (IT)
Day -15
Day 0
Day 1
Day 4
C
TMP489
ND552
Probability of Survival
Days after CAR T injection
CAR-IL9R 0.03E6 + Ad-hIL9
CAR-IL9R 0.03E6 + isotype ctrl
CAR-IL9R 0.03E6 + α-mIL9
Ad-hIL9
ns
***
****
**
*
****
D
TMP489
ND552
Ad-hIL9
CAR-IL9R 0.03E6 + α-mIL9
CAR-IL9R 0.03E6 + isotype ctrl
CAR-IL9R 0.03E6 + Ad-hIL9
Normalized tumor volume
Normalized tumor size
Days after CAR T injection
E
TMP489
Week 2
Week 5
Week 7
cell count/µl
CAR-IL9R T 0.03E6 + Ad-hIL9
CAR-IL9R T 0.03E6 + isotype ctrl
CAR-IL9R 0.03E6 + α-mIL9

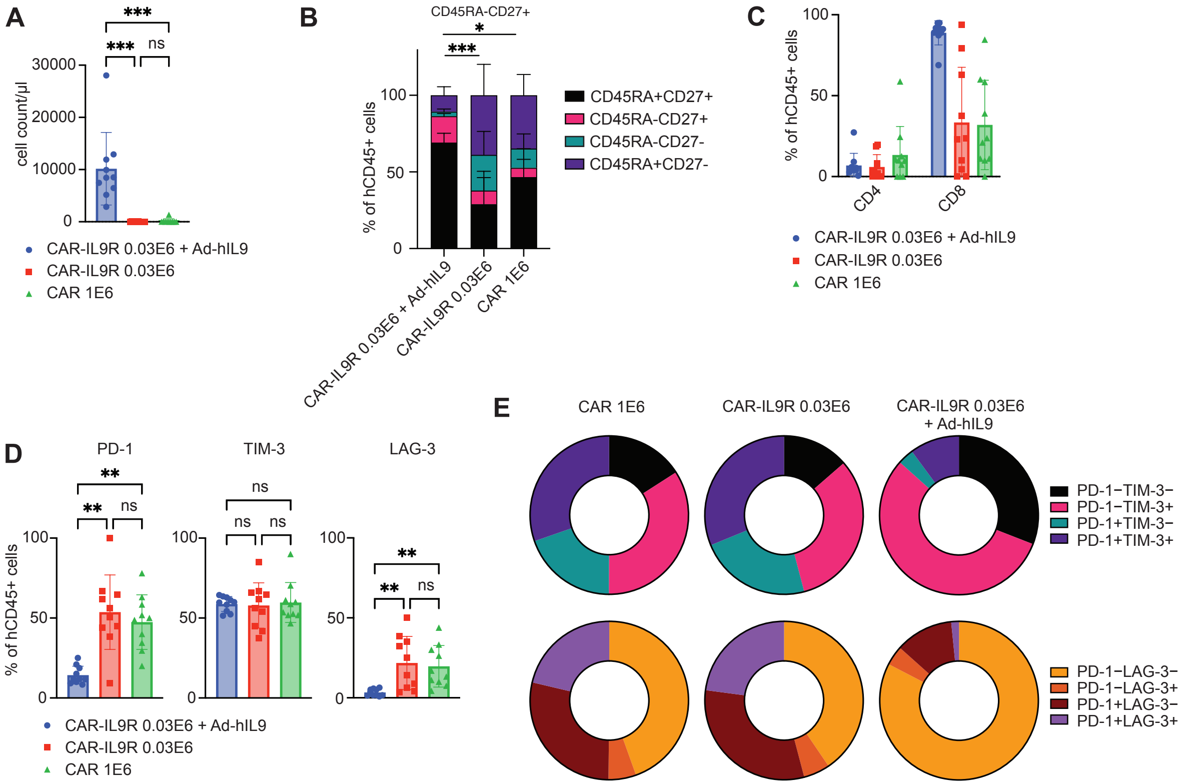

A
***
***
ns
cell count/µl
30000
20000
10000
0
CAR-IL9R 0.03E6 + Ad-hIL9
CAR-IL9R 0.03E6
CAR 1E6
B
CD45RA-CD27+
*
***
% of hCD45+ cells
100
50
0
CAR-IL9R 0.03E6 + Ad-hIL9
CAR-IL9R 0.03E6
CAR 1E6
CD45RA+CD27+
CD45RA-CD27+
CD45RA-CD27-
CD45RA+CD27-
C
% of hCD45+ cells
100
50
0
CD4
CD8
CAR-IL9R 0.03E6 + Ad-hIL9
CAR-IL9R 0.03E6
CAR 1E6
D
PD-1
TIM-3
LAG-3
**
**
ns
ns
ns
ns
**
**
ns
% of hCD45+ cells
100
50
0
CAR-IL9R 0.03E6 + Ad-hIL9
CAR-IL9R 0.03E6
CAR 1E6
E
CAR 1E6
CAR-IL9R 0.03E6
CAR-IL9R 0.03E6 + Ad-hIL9
PD-1−TIM-3−
PD-1−TIM-3+
PD-1+TIM-3−
PD-1+TIM-3+
PD-1−LAG-3−
PD-1−LAG-3+
PD-1+LAG-3−
PD-1+LAG-3+

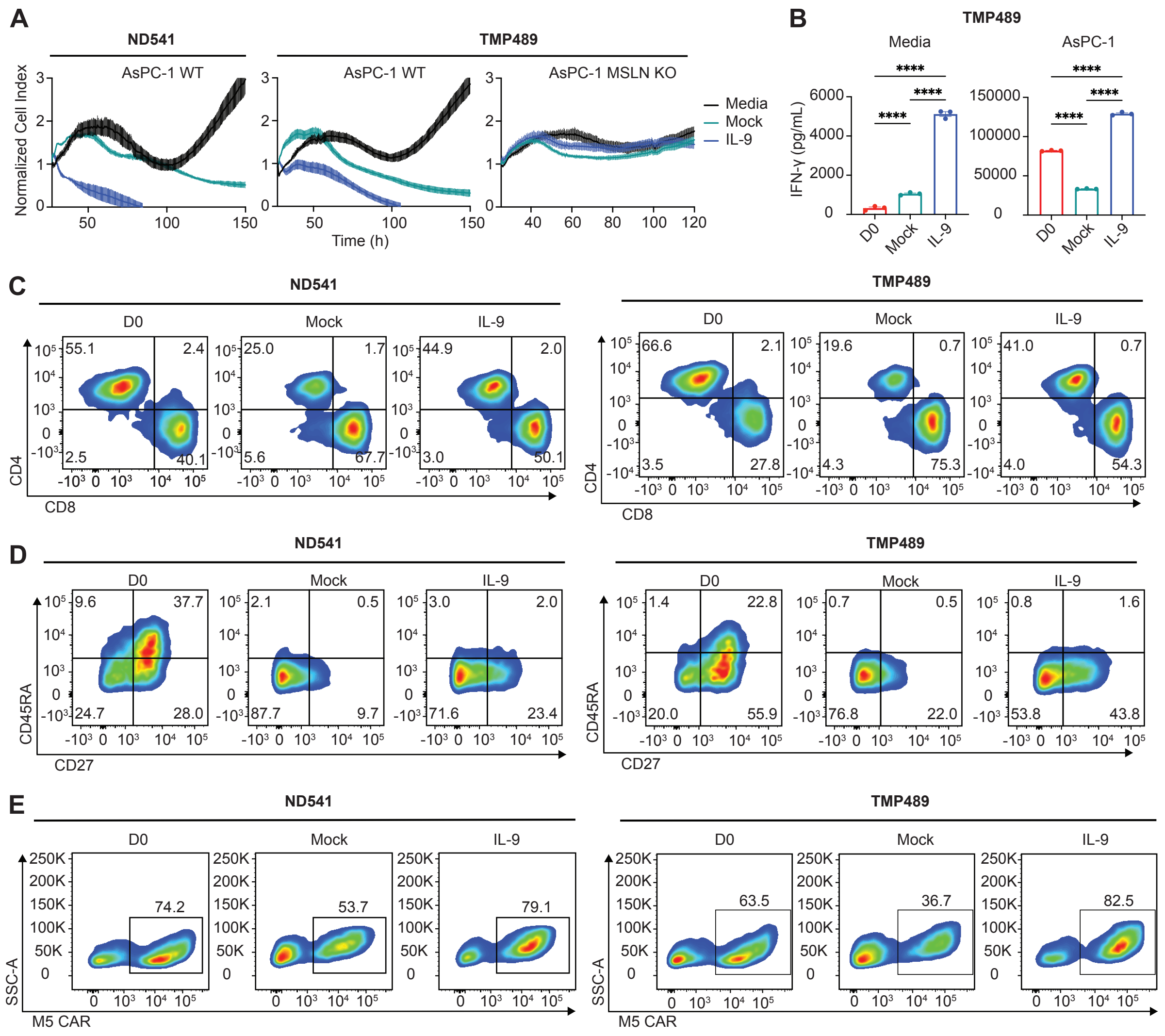

A
ND541
TMP489
AsPC-1 WT
AsPC-1 WT
AsPC-1 MSLN KO
Normalized Cell Index
Time (h)
Media
Mock
IL-9
B
TMP489
Media
AsPC-1
IFN-γ (pg/mL)
****
D0
Mock
IL-9
C
ND541
TMP489
D0
Mock
IL-9
CD4
CD8
55.1
2.4
2.5
40.1
25.0
1.7
5.6
67.7
44.9
2.0
3.0
50.1
66.6
2.1
3.5
27.8
19.6
0.7
4.3
75.3
41.0
0.7
4.0
54.3
D
ND541
TMP489
CD45RA
CD27
9.6
37.7
24.7
28.0
2.1
0.5
87.7
9.7
3.0
2.0
71.6
23.4
1.4
22.8
20.0
55.9
0.7
0.5
76.8
22.0
0.8
1.6
53.8
43.8
E
ND541
TMP489
SSC-A
M5 CAR
74.2
53.7
79.1
63.5
36.7
82.5

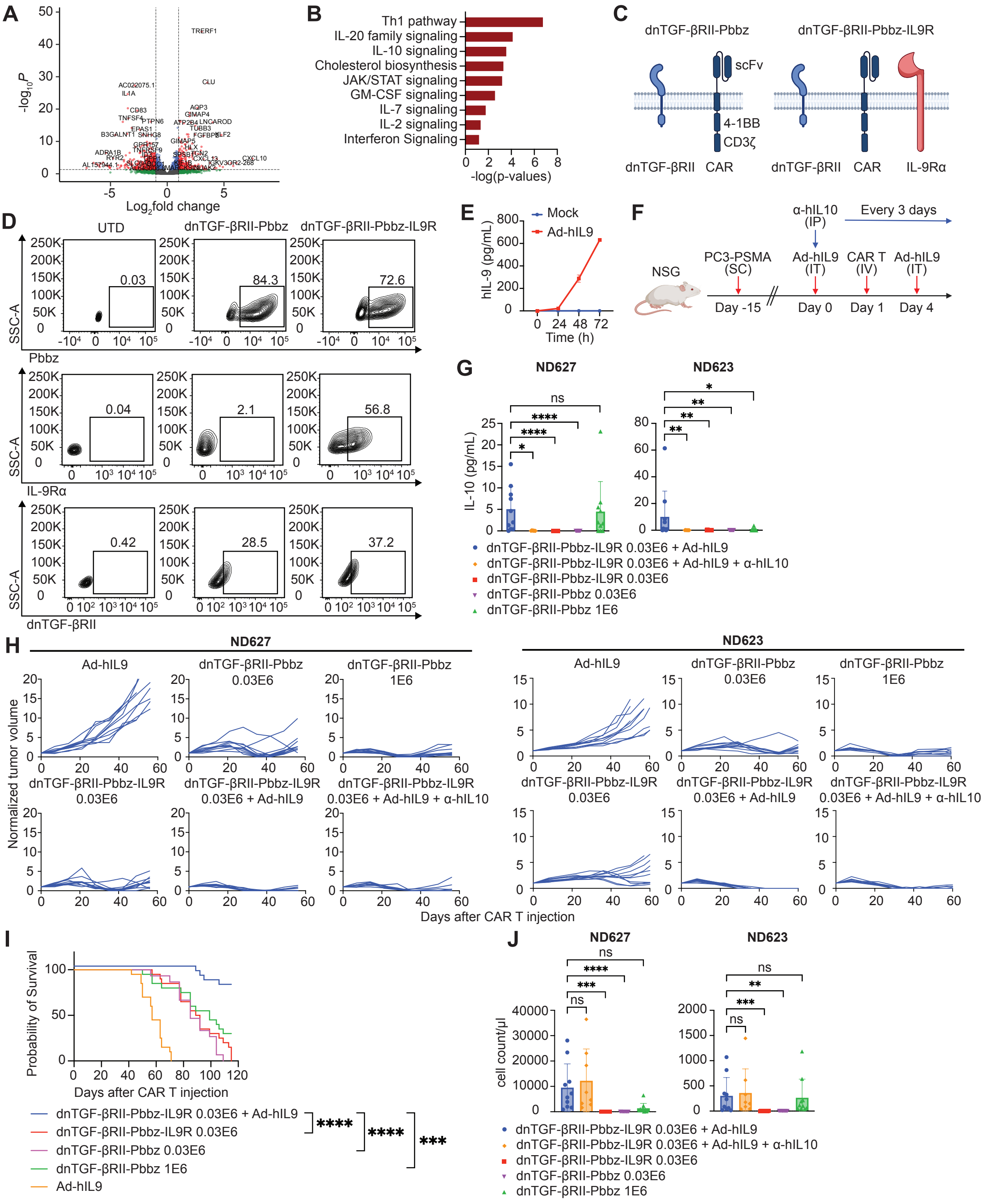

A
B
Th1 pathway
IL-20 family signaling
IL-10 signaling
Cholesterol biosynthesis
JAK/STAT signaling
GM-CSF signaling
IL-7 signaling
IL-2 signaling
Interferon Signaling
-log(p-values)
C
dnTGF-βRII-Pbbz
dnTGF-βRII-Pbbz-IL9R
scFv
4-1BB
CD3ζ
dnTGF-βRII
CAR
IL-9Rα
D
UTD
Pbbz
IL-9Rα
dnTGF-βRII
SSC-A
E
Mock
Ad-hIL9
hIL-9 (pg/mL)
Time (h)
F
NSG
PC3-PSMA (SC)
α-hIL10 (IP)
Every 3 days
Ad-hIL9 (IT)
CAR T (IV)
Day -15
Day 0
Day 1
Day 4
G
ND627
ND623
IL-10 (pg/mL)
dnTGF-βRII-Pbbz-IL9R 0.03E6 + Ad-hIL9
dnTGF-βRII-Pbbz-IL9R 0.03E6 + Ad-hIL9 + α-hIL10
dnTGF-βRII-Pbbz-IL9R 0.03E6
dnTGF-βRII-Pbbz 0.03E6
dnTGF-βRII-Pbbz 1E6
H
Normalized tumor volume
Days after CAR T injection
I
Probability of Survival
Days after CAR T injection
Ad-hIL9
J
cell count/μl

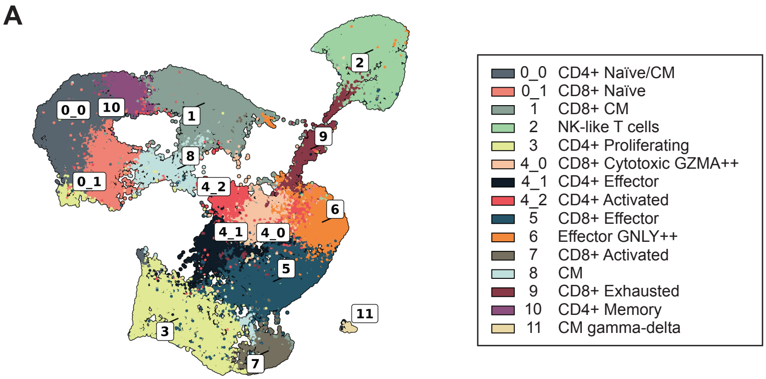
A
0_0
10
1
2
9
8
0_1
4_2
6
4_1
4_0
5
11
3
7
0_0 CD4+ Naïve/CM
0_1 CD8+ Naïve
1 CD8+ CM
2 NK-like T cells
3 CD4+ Proliferating
4_0 CD8+ Cytotoxic GZMA++
4_1 CD4+ Effector
4_2 CD4+ Activated
5 CD8+ Effector
6 Effector GNLY++
7 CD8+ Activated
8 CM
9 CD8+ Exhausted
10 CD4+ Memory
11 CM gamma-delta

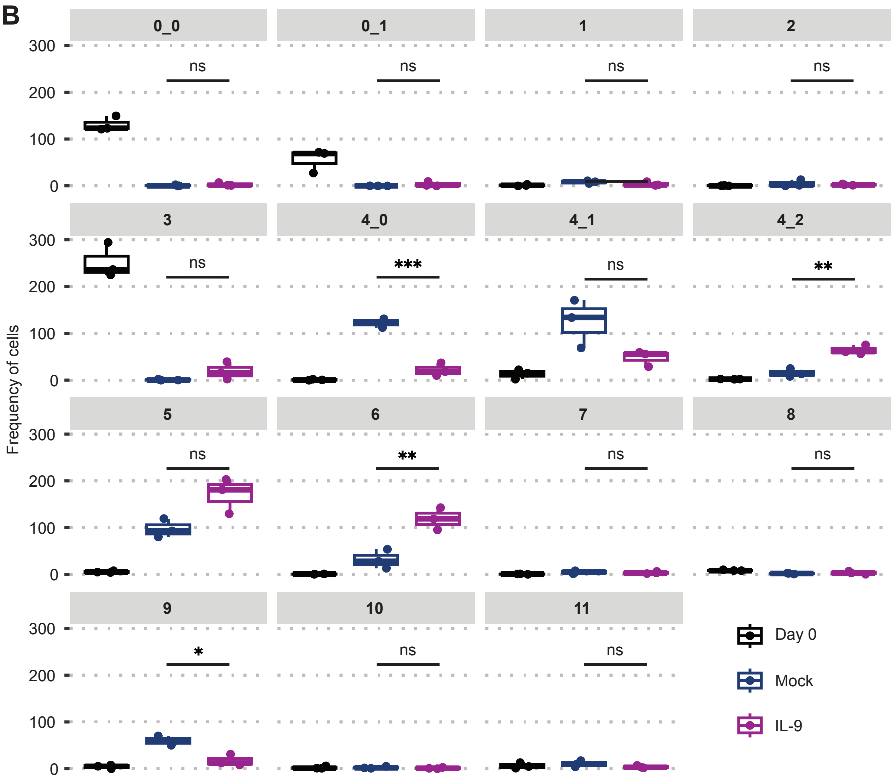
B
0_0
0_1
1
2
3
4_0
4_1
4_2
5
6
7
8
9
10
11
ns
***
**
*
Frequency of cells
300
200
100
0
Day 0
Mock
IL-9

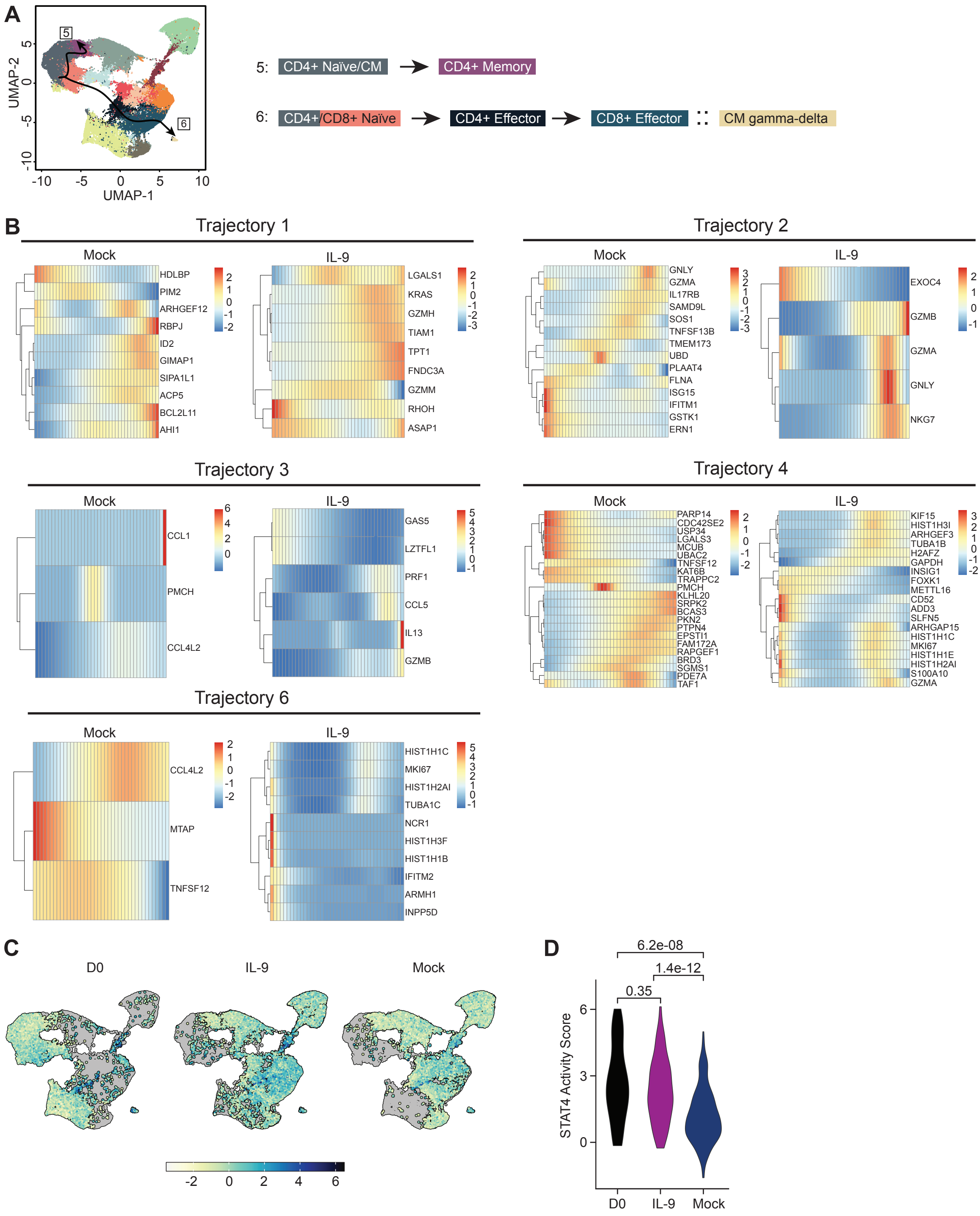
A
UMAP-2
UMAP-1
5: CD4+ Naïve/CM → CD4+ Memory
6: CD4+/CD8+ Naïve → CD4+ Effector → CD8+ Effector :: CM gamma-delta
B
Trajectory 1
Mock
HDLBP
PIM2
ARHGEF12
RBPJ
ID2
GIMAP1
SIPA1L1
ACP5
BCL2L11
AHI1
IL-9
LGALS1
KRAS
GZMH
TIAM1
TPT1
FNDC3A
GZMM
RHOH
ASAP1
Trajectory 2
Mock
GNLY
GZMA
IL17RB
SAMD9L
SOS1
TNFSF13B
TMEM173
UBD
PLAAT4
FLNA
ISG15
IFITM1
GSTK1
ERN1
IL-9
EXOC4
GZMB
GZMA
GNLY
NKG7
Trajectory 3
Mock
CCL1
PMCH
CCL4L2
IL-9
GAS5
LZTFL1
PRF1
CCL5
IL13
GZMB
Trajectory 4
Mock
PARP14
CDC42SE2
USP34
LGALS3
MCUB
UBAC2
TNFSF12
KAT6B
TRAPPC2
PMCH
KLHL20
SRPK2
BCAS3
PKN2
PTPN4
EPSTI1
FAM172A
RAPGEF1
BRD3
SGMS1
PDE7A
TAF1
IL-9
KIF15
HIST1H3I
ARHGEF3
TUBA1B
H2AFZ
GAPDH
INSIG1
FOXK1
METTL16
CD52
ADD3
SLFN5
ARHGAP15
HIST1H1C
MKI67
HIST1H1E
HIST1H2AI
S100A10
GZMA
Trajectory 6
Mock
CCL4L2
MTAP
TNFSF12
IL-9
HIST1H1C
MKI67
HIST1H2AI
TUBA1C
NCR1
HIST1H3F
HIST1H1B
IFITM2
ARMH1
INPP5D
C
D0
IL-9
Mock
D
6.2e-08
1.4e-12
0.35
STAT4 Activity Score
D0
IL-9
Mock
